# Genome sequence of *Hydrangea macrophylla* and its application in analysis of the double flower phenotype

**DOI:** 10.1101/2020.06.14.151431

**Authors:** K Nashima, K Shirasawa, A Ghelfi, H Hirakawa, S Isobe, T Suyama, T Wada, T Kurokura, T Uemachi, M Azuma, M Akutsu, M Kodama, Y Nakazawa, K Namai

## Abstract

Owing to its high ornamental value, the double flower phenotype of hydrangea (*Hydrangea macrophylla*) is one of its most important traits. In this study, genome sequence information was obtained to explore effective DNA markers and the causative genes for double flower production in hydrangea. Single molecule real-time sequencing data followed by a HiC analysis was employed. The resultant haplotype-phased sequences consisted of 3,779 sequences (2.256 Gb in length and N50 of 1.5 Mb), and 18 pseudomolecules comprising 1.08 Gb scaffold sequences along with a high-density SNP genetic linkage map. Using the genome sequence data obtained from two breeding populations, the SNPs linked to double flower loci (*D*_*jo*_ and *D*_*su*_), were discovered for each breeding population. DNA markers J01 linked to *D*_*jo*_ and S01 linked to *D*_*su*_ were developed, and these could be used successfully to distinguish the recessive double flower allele for each locus respectively. The *LEAFY* gene was suggested as the causative gene for *D*_*su,*_ since frameshift was specifically observed in double flower accession with *d*_*su*_. The genome information obtained in this study will facilitate a wide range of genomic studies on hydrangea in the future.

## 1. Introduction

*Hydrangea macrophylla* (Thunb.) Ser., commonly known as hydrangea, originated in Japan, and since it is the place of origin, there are rich genetic resources for this plant in Japan. Wild hydrangea accessions with superior characteristics have been bred to create attractive cultivars, and it has a long history of use as an ornamental garden plant in temperate regions. There are both decorative and non-decorative flowers in an inflorescence. Decorative flowers have large ornamental sepals that attract pollinators, whereas non-decorative flowers have inconspicuous perianths that instead play a major role in seed production^1–3^. In hydrangea, there are two types of decorative flower phenotype: single flower and double flower. Single flowers generally have four petaloid sepals per decorative flower, while this number in double flowers is approximately fourteen. Double flowers do not have stamens or petals^4^. Therefore, petals and stamens would be converted to petaloid sepals since number of petaloid sepals are increased and stamens and petals are lost. Because of their high ornamental value, producing double flower is an important breeding target in hydrangea cultivation.

To obtain double flower progenies, the double flower cultivars ‘Sumidanohanabi’ (Figure 1A) and ‘Jogasaki’ (Figure 1B) were crossbred in Japan^4^. Previous studies have suggested that double flower phenotype is a recessive characteristic controlled by a single major gene^4,5^. Suyama et al.^4^ found that crosses between the progeny of ‘Sumidanohanabi’ and the progeny of ‘Jogasaki’ produced only single flower descendants. Thus, it was also suggested that genes controlling the double flower phenotype are different^4^. While Suyama et al.^4^ suggested that a single locus with different double flower alleles controls the phenotype, Waki et al.^5^ speculated that two different loci control double flower production individually. Therefore, it is not clear whether a single locus or two loci control the phenotype. We term the double flower locus *D*_*su*_ as the locus controlling the double flower phenotype of ‘Sumidanohanabi’ and the double flower locus *D*_*jo*_ as the locus controlling the double flower phenotype of ‘Jogasaki.’ Waki et al.^5^ identified *D*_*su*_ on the genetic linkage map. They also found that the DNA marker STAB045 was the nearest marker to *D*_*su*_, and that STAB045 could help in distinguishing flower phenotype with a 98.6% fitting ratio^5^. Contrarily, *D*_*jo*_ has not been identified, and the DNA marker linked to *D*_*jo*_ has not been developed. It is still not known whether *D*_*jo*_ and *D*_*su*_ are at the same loci.

**Figure 1.**
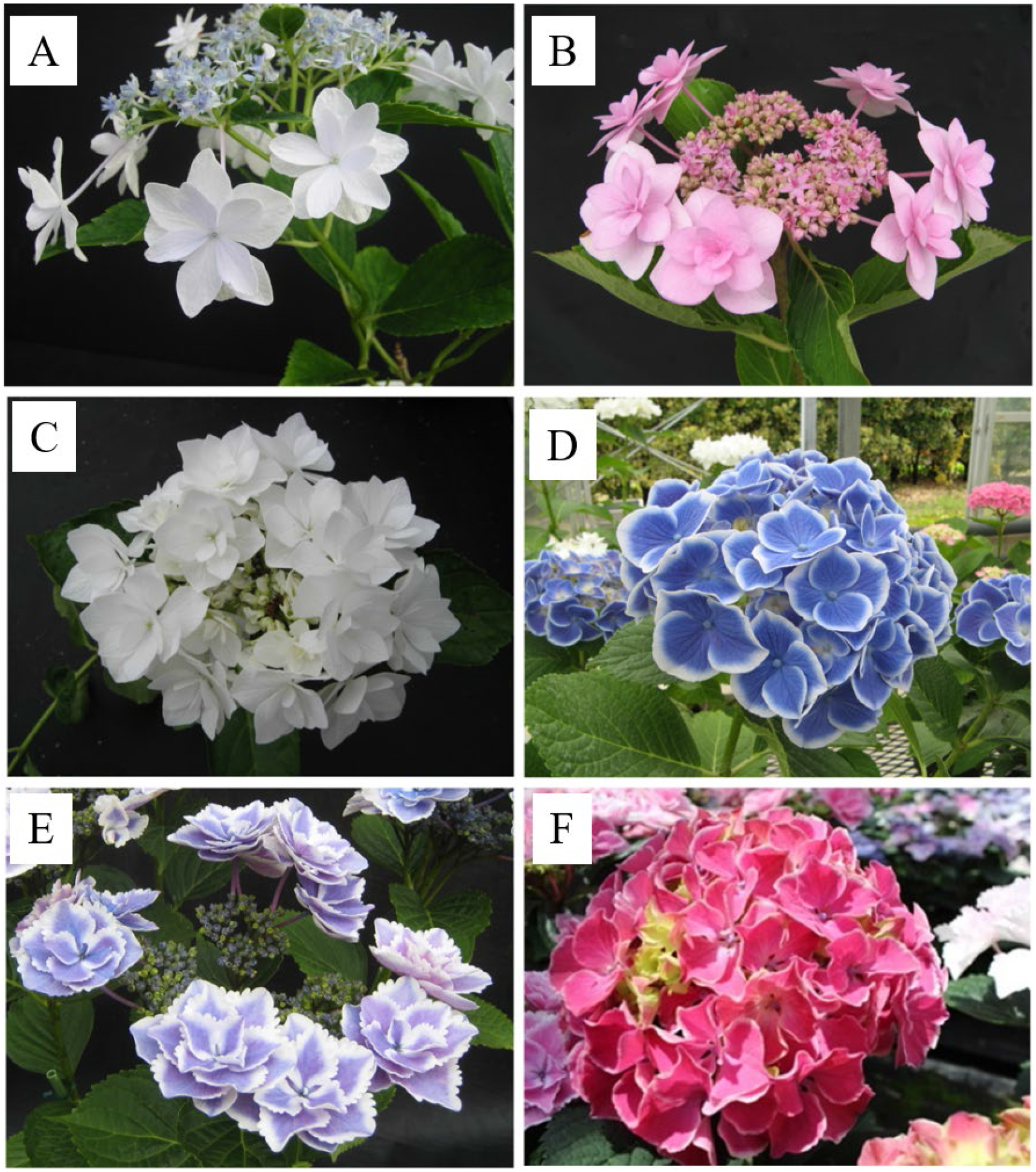
Flower phenotypes of hydrangea accessions A: ‘Sumidanohanabi’ (double flower). B: ‘Jogasaki’ (double flower). C: ‘Posy Bouquet Grace’ (double flower). D: ‘Blue Picotee Manaslu’ (single flower). E: ‘Kirakiraboshi’ (double flower). F: ‘Frau Yoshimi’ (single flower).

The mechanisms and genes controlling double flower phenotype in hydrangea have not been clarified. Waki et al.^5^ hypothesized that the mutation of C-class genes could be associated with the double flower phenotype of ‘Sumidanohanabi’, since the C-class gene mutant of *Arabidopsis thaliana* and C-class gene-repressed petunias produce double flowers^6^. However, the double flower phenotype of hydrangea is morphologically different from that of *A. thaliana* and petunia—petals and stamens would be converted to petaloid sepals, while stamens converted to petals in *A. thaiana* and petunia. This suggests that the genes controlling double flower production in hydrangea are different from corresponding genes in other plant species. Identification of the genes controlling double flower production in hydrangea could reveal novel regulatory mechanisms of flower development.

Genomic information is essential for DNA marker development and identification of genes controlling specific phenotypes. However, no reference genome sequence is publicly available for hydrangea so far. Although a genome assembly of hydrangea (1.6 Gb) using only short-read data has been reported^7^, the resultant assembly is so fragmented that it comprises 1,519,429 contigs with an N50 size of 2,447 bp and has not been disclosed. Improved, advanced long-read technologies and bioinformatics methods would make it possible to determine the sequences of complex genomes. An assembly strategy for single molecule real-time sequencing data followed by a HiC analysis has been developed to generate haplotype-phased sequences in heterozygous regions of diploid genomes^8^. Genome sequences at the chromosome level could be obtained with a HiC clustering analysis^9^ as well as with a genetic linkage analysis^10^. Such genomic sequence will provide basic information to identify genes and DNA markers of interest, and to discover allelic sequence variations. In this study, we constructed the genomic DNA sequence, obtained SNPs information, and performed gene prediction. We also developed DNA markers linked to *D*_*jo*_ using SNP information obtained by double digest restriction site associated DNA sequence (ddRAD-Seq) analysis of breeding population 12GM1, which segregated double flower phenotypes of *D_jo_*. In addition, we attempted to identify the causative genes for *D*_*jo*_ and *D*_*su*_.

## 2. Materials and Methods

### 2.1. De novo assembly of the hydrangea genome

For genomic DNA sequencing, *H. macrophylla* ‘Aogashima-1,’ collected from Aogashima island of the Izu Islands in Tokyo Prefecture, Japan, was used. Genomic DNA was extracted from the young leaves with Genomic-Tip (Qiagen, Hilden, Germany). First, we constructed a sequencing library (insert size of 500 bp) with TruSeq DNA PCR-Free Library Prep Kit (Illumina, San Diego, CA, USA) to sequence on HiSeqX (Illumina). The size of the 'Aogashima-1' genome was estimated using Jellyfish v2.1.4^11^. After removing adapter sequences and trimming low-quality reads, high-quality reads were assembled using Platanus^12^. The resultant sequences were designated HMA_r0.1. Completeness of the assembly was assessed with sets of BUSCO v.1.1b^13^.

Next, a SMRT library was constructed with SMRTbell Express Template Prep Kit 2.0 (PacBio, Menlo Park, CA, USA) in accordance with the manufacture’s protocol and sequenced with SMRT Cell v2.1 on a Sequel System (PacBio). The sequence reads were assembled using FALCON v.1.8.8^14^ to generate primary contig sequences and to associate contigs representing alternative alleles. Haplotype - resolved assemblies (i.e. haplotigs) were generated using FALCON-Unzip v.1.8.8^14^. Potential sequence errors in the contigs were corrected twice with ARROW v.2.2.1 implemented in SMRT Link v.5.0 (PacBio) followed by one polishing with Pilon^15^. Subsequently, a HiC library was constructed with Proximo Hi-C (Plant) Kit (Phase Genomics, Seattle, WA, USA) and sequenced on HiSeqX (Illumina). After removing adapter sequences and trimming low-quality reads, high-quality HiC reads were used to generate two haplotype-phased sequences from the primary contigs and haplotig sequences with FALCON-Phase^8^.

To validate the accuracy of the sequences, we developed a genetic map based on SNPs, which were from a ddRAD-Seq analysis on an F2 mapping population (n = 147), namely 12GM1, maintained at the Fukuoka Agriculture and Forestry Research Center, Japan. The 12GM1 population was generated from a cross between ‘Posy Bouquet Grace’ (Figure 1C) and ‘Blue Picotee Manaslu’ (Figure 1D). Genomic DNA was extracted from the leaves with DNeasy Plant Mini Kit (Qiagen). A ddRAD-Seq library was constructed as described in Shirasawa et al.^16^ and sequenced with HiSeq4000. Sequence reads were processed as described by Shirasawa et al.^16^ and mapped on the HMA_r1.2 as a reference. From the mapping alignment, high-confidence biallelic SNPs were obtained with the following filtering options: --minDP 5 --minQ 10 --max-missing 0.5. The genetic map was constructed with Lep-Map3^17^.

Potential mis-jointed points in the phase 0 and 1 sequences of HMA_r1.2 were cut and re-joined, based on the marker order in the genetic map, for which we employed ALLMAPS^18^. The resultant sequences were named HMA_r1.3.pmol, as two haplotype-phased pseudomolecule sequences of the ‘Aogashima-1’ genome. Sequences that were unassigned to the genetic map were connected and termed chromosome 0.

### 2.2 Gene prediction

For gene prediction, we performed Iso-Seq analysis. Total RNA was extracted from 12 samples of ‘Aogashima-1’: flower buds (2 stages); decorative flowers (2 stages); colored and colorless non-decorative flowers; fruits; shoots; roots; buds, and one-day light-intercepted leaves and buds. In addition, the 29 samples listed in Supplementary Table S1 were included. Iso-Seq libraries were prepared with the manufacture’s Iso-Seq Express Template Preparation protocol, and sequenced on a Sequel System (PacBio). The raw reads obtained were treated with ISO-Seq3 pipeline, implemented in SMRT Link v.5.0 (PacBio) to generate full-length, high-quality consensus isoforms. In parallel, RNA-Seq data was also obtained from the 16 samples listed in Supplementary Table S1. Total RNA extracted from the samples was converted into cDNA and sequenced on HiSeq2000, Hiseq2500 (Illumina), and NovaSeq6000 (Illumina). The Iso-Seq isoform sequences and the RNA-Seq short-reads were employed for gene prediction.

To identify putative protein-encoding genes in the genome assemblies, ab-initio-, evidence-, and homology-based gene prediction methods were used. For this prediction, unigene sets generated from 1) the Iso-Seq isoforms; 2) de novo assembly of the RNA-Seq short-reads with Trinity-v2.4.0^19^; peptide sequences predicted from the genomes of *Arabidopsis thaliana*, *Arachis hypogaea*, *Cannabis sativa*, *Capsicum annuum*, *Cucumis sativus*, *Populus trichocarpa*, and *Quercus lobata*; and *ab-initio* genes, were predicted with Augustus-v3.3.1^20^. The unigene sequences were aligned onto the genome assembly with BLAT^21^ and genome positions of the genes were listed in general feature format version 3 with blat2gff.pl (https://github.com/vikas0633/perl/blob/master/blat2gff.pl). Gene annotation was performed with Hayai-annotation Plants^22^. Completeness of the gene prediction was assessed with sets of BUSCO v4.0.6^13^.

### 2.3 Detection of SNPs linked to the double flower phenotype

For identification of SNPs linked to double flower loci *D*_*jo*_ and *D*_*su*_, ddRAD-Seq data analysis was performed. ddRAD-Seq data of the 12GM1 population described above was used to identify *D*_*jo*_. For identification of SNPs linked to double flower locus *D*_*su*_, KF population^5^—93 F2 specimens of ‘Kirakiraboshi’ (Figure 1E) and ‘Frau Yoshimi’ (Figure 1F)—were used for ddRAD-Seq analysis. The KF population was maintained at Tochigi Prefectural Agricultural Experimental Station, Japan. ddRAD-Seq analysis of the KF population was performed using the same method used for the 12GM1 population.

ddRAD-Seq data of the 12GM1 and KF populations were processed as follows: Low-quality sequences were removed and adapters were trimmed using Trimmomatic-0.36^23^ (LEADING:10, TRAILING:10, SLIDINGWINDOW:4:15, MINLEN:51). BWA-MEM (version 0.7.15-r1140) was used for mapping onto genome sequence. The resultant sequence alignment/map format (SAM) files were converted to binary sequence alignment/map format files and subjected to SNP calling using the mpileup option of SAMtools^24^ (version 1.4.1) and the view option of BCFtools (parameter -vcg). If the DP of called SNP in individuals was under 5%, the genotype was treated as missing. SNPs with 5% or more of missing genotype were filtered out. Each SNP was evaluated, fitting ratios with the flower phenotype.

### 2.4 DNA marker development and analysis for D_jo_

A CAPS marker was designed based on SNP (Scaffold:0008F-2, position: 780104) that was completely linked to the double flower locus *D*_*jo*_. Primers were designed using Primer3^25^ under conditions with product size ranging from 150 to 350 bp, primer size from 18 to 27 bp, and primer TM from 57 to 63°C. Primer sequences of the designed CAPS marker named J01 were: Forward: 5′-CTGGCAGATTCCTCCTGAC-3′ and Reverse: 5′-TATTTCCTTGGGGAGGCTCT-3′. PCR assays were done in a total volume of 10 μL, containing 5 μL of GoTaq Master Mix (Promega, Mdison, WI, USA), 1 mM each of forward and reverse primer, and 5 ng of template DNA. The PCR conditions were 94°C for 2 min, 35 cycles of denaturation at 94°C for 1 min, annealing at 55°C for 1 min, and extension at 72°C for 1 min; and a final extension step at 72°C for 3 min. Then, restriction enzyme assay was done in a total volume of 10 μL, containing 5 μL of PCR product, ten units of restriction enzyme TaqI (New England Biolabs, Ipswich, MA, USA), and 1 μL of cut smart buffer. Restriction enzyme assay was performed at 65°C for 3 h. The restriction assay product was stained with 1x GRRED (Biocraft, Tokyo, Japan) and separated in 1.5% (w/v) agarose gel in TAE buffer. Designed CAPS marker J01 was applied to the 12GM1 population, 14GT77 population (64 F2 specimens of ‘Posy Bouquet Grace’ × ‘Chibori’) and the 15IJP1 population (98 F1 specimens of ‘Izunohana’ × 03JP1) that segregate the double flower locus *D*_*jo*_.

### 2.5 Resequencing and comparison of LEAFY gene sequence and DNA marker development

To compare sequences, resequencing of genomic DNA was performed for accessions of ‘Kirakiraboshi,’ ‘Frau Yoshimi,’ ‘Posy Bouquet Grace,’ and ‘Blue Picotee Manaslu.’ Sequencing libraries (insert size of 500 bp) for the four lines were constructed with TruSeq DNA PCR-Free Library Prep Kit (Illumina) to sequence on a HiSeqX (Illumina). From the sequence reads obtained, low-quality bases were deleted with PRINSEQ v.0.20.4^26^ and adaptor sequences were trimmed with fastx clipper (parameter, -a AGATCGGAAGAGC) in FASTX-Toolkit v.0.0.13 (http://hannonlab.cshl.edu/fastx_toolkit). High-quality reads were aligned on the HMA_r1.2 with Bowtie2^27^ v.2.2.3 to detect sequence variant candidates by with the mpileup command in SAMtools v.0.1.19^24^. High-confidence variants were selected using VCFtools^29^ v.0.1.12b with parameters of -- minDP 10, --maxDP 100, --minQ 999, --max-missing 1.

For comparison of *LEAFY (LFY)* sequence in ‘Kirakiraboshi,’ ‘Frau Yoshimi,’ ‘Posy Bouquet Grace,’ and ‘Blue Picotee Manaslu,’ BLAST analysis using genomic sequence of *LFY* (Scaffold 0577F, position 678200-684639) as query, and genomic DNA sequence of each cultivar as database, was performed to confirm detected sequence variants. These data analyses were performed using CLC main workbench (Qiagen). INDEL marker S01 that amplifies the second intron of *LFY*, was designed by visual inspection (Forward: 5′-CATCATTAATAGTGGTGACAG-3′, Reverse: 5′-CACACATGAATTAGTAGCTC-3′). The PCR conditions were 94°C for 2 min, 35 cycles of denaturation at 94°C for 1 min, annealing at 55°C for 1 min, extension at 72°C for 1 min; and a final extension step at 72°C for 3 min. The PCR product was stained with 1x GRRED (Biocraft) and separated in 2.5% (w/v) agarose gel in TAE buffer.

### 2.6 Cloning and sequence determination of LFY gene of ‘Kirakiraboshi’ and ‘Frau Yoshimi’

Total RNA was isolated from the flower buds of ‘Kirakiraboshi,’ and ‘Frau Yoshimi’ using RNAiso Plus (TaKaRa, Japan), and reverse transcribed using PrimeScriptⅡ 1st strand cDNA Synthesis Kit (TaKaRa, Japan). The sequence of the *LFY* gene was amplified by PCR in 50-µL reaction mixture by using TaKaRa Ex Taq Hot Start Version (TaKaRa Bio, Shiga, Japan) and the *LFY* specific primer (Forward: 5′-ATGGCTCCACTACCTCCACC-3′ and Reverse: 5′-CTAACACCCTCTAAAAGCAG-3′). These PCR products were purified, and inserted into a pMD20-T vector using the Mighty TA-cloning kit (TaKaRa Bio). The sequence of *LFY* coding sequence (CDS) in pMD20-T vector was analyzed by 3130xl DNA sequencer (Applied Biosystems, Foster City, CA, USA). Sequence alignments were obtained by using CLC main workbench (Qiagen).

### 2.7 DNA marker assessment across hydrangea accessions

For assessment of DNA markers for the double flower phenotype, 35 *H. macrophylla* accessions were used. Genotyping for J01 was performed as described above. Genotyping for S01 was performed by fragment analysis as follows. PCR amplification was performed in a 10- μL reaction mixture containing 5 μL of GoTaq Master Mix (Promega), 5 pmol FAM-labeled universal primer (5′ - FAM-gctacggactgacctcggac-3′), 2.5 pmol forward primer with universal adapter sequence (5′-gctacggactgacctcggacCATCATTAATAGTGGTGACAG-3′), 5 pmol reverse primer, and 5 ng of template DNA. DNA was amplified in 35 cycles of 94°C for 1 min, 55°C for 1 min, and 72°C for 2 min; and a final extension of 5 min at 72°C. The amplified PCR products were separated and detected in a PRISM 3130xl DNA sequencer (Applied Biosystems, USA). The sizes of the amplified bands were scored against internal-standard DNA (400HD-ROX, Applied Biosystems, USA) by GeneMapper software (Applied Biosystems, USA).

## 3. Results and Discussion

### 3.1 Draft genome assembly with long-read and HiC technologies

The size of the hydrangea genome was estimated by k-mer-distribution analysis with the short-read of 132.3 Gb data. The resultant distribution pattern indicated two peaks, representing homozygous (left peak) and heterozygous (right peak) genomes, respectively (Figure 2). The haploid genome of hydrangea was estimated to be 2.2 Gb in size. The short reads were assembled into 612,846 scaffold sequences. The total length of the resultant scaffolds, i.e. HMA_r0.1, was 1.7 Gb with an N50 length of 9.1 kb (Supplementary Table S2). Only 72.2% of complete single copy orthologues in plant genomes were identified in a BUSCO analysis (Supplementary Table S2).

**Figure 2.**
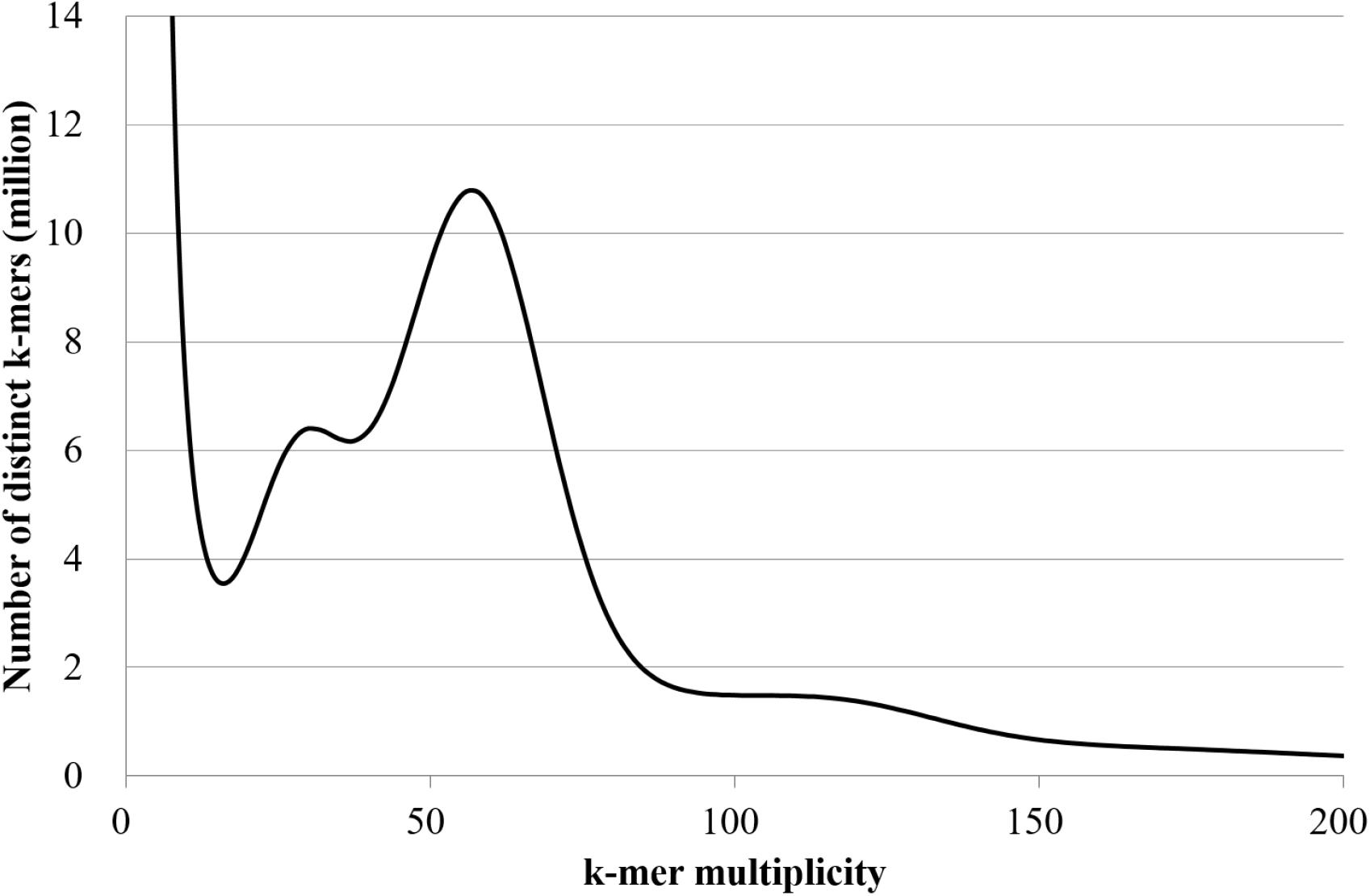
Genome size estimation for the hydrangea line ‘Aogashima-1’ with the distribution of the number of distinct *k*-mers (*k*=17), with the given multiplicity values.

Next, we employed long sequence technology to extend the sequence contiguity and to improve the genome coverage. A total of 106.9 Gb of reads (49.4×) with an N50 read length of 28.8 kb was obtained from 14 SMRT Cells. The long-reads were assembled, followed by sequence error corrections into 15,791 contigs consisting of 3,779 primary contigs (2.178 Gb in length and N50 of 1.4 Mb), and 12,012 haplotig sequences (1.436 Gb in length and N50 of 184 kb). To obtain two haplotype-phased complete-length sequences, 697 M reads of HiC data (105.3 Gb) were obtained and subjected to FALCON-Phase. The resultant haplotype-phased sequences consisted of 3,779 sequences (2.256 Gb in length and N50 of 1.5 Mb) for “phase 0,” and 3,779 sequences (2.227 Gb in length, and N50 of 1.4 Mb) for “phase 1.”

### 3.2 Pseudomolecule sequences based on genetic mapping

To detect potential errors in the assembly and to assign the contig sequences onto the hydrangea chromosomes, we established an F2 genetic map based on SNPs derived from a ddRAD-Seq technology. Approximately 1.8 million high-quality ddRAD-Seq reads per sample were obtained from the mapping population and mapped to either of the two phased sequences with alignment rates of 88.4% and 88.7%, respectively. A set of SNPs detected from the alignments were classified into 18 groups and ordered to construct two genetic maps for the two phased sequences (2,849.3 cM in length with 3,980 SNPs, and 2,944.5 cM in length with 4,071 SNPs). The nomenclature of the linkage groups was named in accordance with the previous genetic map based on SSRs^5^. The phased sequences were aligned on each genetic map to establish haplotype-phased, chromosome-level pseudomolecule sequences. During this process, one contig was cut due to possible mis-assembly. The resultant sequences for phase 0 had 730 contigs with a total length of 1,078 Mb and the other for phase 1 had 743 contigs spanning 1,076 Mb.

### 3.3. Transcriptome analysis followed by gene prediction

In the Iso-Seq analysis, Circular Consensus Sequence (CCS) reads were generated from the raw sequence reads. The CCS reads were classified in full-length and non-full length reads and the full-length reads were clustered to produce consensus isoforms. In total, 116,634 high-quality isoforms were used for gene prediction. In the RNA-Seq analysis, on the contrary, a total of 80.7 Gb reads were obtained and assembled into 12,265 unigenes. The high-quality isoforms and unigenes together with gene sequences predicted from the *Arabidopsis thaliana*, *Arachis hypogaea*, *Cannabis sativa*, *Capsicum annuum*, *Cucumis sativus*, *Populus trichocarpa*, and *Quercus lobate* genomes were aligned onto the assembly sequence of the hydrangea genome. By adding ab-initio on genes, 32,205 and 32,222 putative protein-encoding genes were predicted from the phase 0 and phase 1 sequences, respectively. This gene set included 91.4% complete BUSCOs. Out of the 10,108 genes, 16,725, and 21,985 were assigned to Gene Ontology slim terms in the biological process, cellular component, and molecular function categories, respectively. Furthermore, 4,271 genes had assigned enzyme commission numbers.

### 3.4 Identification of SNPs tightly linked to double flower phenotype

To identify SNPs tightly linked to the double flower phenotype of ‘Jogasaki,’ ddRAD-Seq analysis was performed on the 12GM1 population, which segregates the double flower phenotype of ‘Jogasaki.’ As a result, 14,006 of SNPs were called by ddRAD-Seq analysis of the 12GM1 population. In this population, the double flower phenotype was expected when the plant was homozygous for the ‘Posy Bouquet Grace’ genotype, and the single flower phenotype was expected when the plant was homozygous for ‘Blue Picotee Manaslu’ or was heterozygous. Each SNP was tested for its fitting rate to this model. As a result, ten SNPs were found to have more than a 95% fitting rate, and six SNPs were completely co-segregated with flower phenotype (Table 1).

**Table 1.** SNPs correlated (fitting rate more than 95%) with double flower phenotype in 12GM1 population

| Scaffold | Position at Phase 0 | Sequence variant |  | Fitting rate (%) | Frequency of double flower phenotype (double flower/all) |  |  |
| --- | --- | --- | --- | --- | --- | --- | --- |
|  |  | Posy Bouquet Grace | Blue Picotee manasulu |  | Homozygous of 'Posy Bouquet Grace' | Heterozygous | Homozygous of 'Blue Picotee Manasulu' |
| 0008F-2 | 3250598 | A | G | 100 | 37/37 | 0/61 | 0/47 |
| 0008F-2 | 3250523 | A | C | 100 | 37/37 | 0/61 | 0/47 |
| 0008F-2 | 780104 | C | A | 100 | 37/37 | 0/60 | 0/48 |
| 0259F | 404610 | T | A | 100 | 37/37 | 0/60 | 0/48 |
| 1207F | 365533 | C | T | 100 | 38/38 | 0/61 | 0/48 |
| 1207F | 372121 | C | A | 100 | 38/38 | 0/61 | 0/47 |
| 0012F | 1318350 | T | C | 97.9 | 37/39 | 1/59 | 0/48 |
| 0437F | 170787 | G | A | 97.9 | 36/37 | 1/60 | 1/49 |
| 0437F | 180821 | A | G | 97.9 | 36/37 | 1/60 | 1/49 |
| 0994F | 216439 | C | T | 97.9 | 36/37 | 1/60 | 1/49 |

CAPS marker J01 was developed based on SNP at scaffold 0008F-2_780104. J01 CAPS marker amplified 167 bp of fragment by PCR, and digestion with Taq I restriction enzyme generated 50 bp and 117 bp fragments in the double flower allele (Figure 3). J01 marker was fitted with flower phenotype at 99.3% in the 15IJP1 and 14GT77 populations, which segregated the double flower phenotype of ‘Jogasaki’ (Supplementary Table S3, S4). This indicated that J01 marker was tightly linked to the *D*_*jo*_ locus. Thus, *D*_*jo*_ is suggested to be located adjacent to J01, which is located at position 46,326,384 in CHR17, (Figure 4).

**Figure 3.**
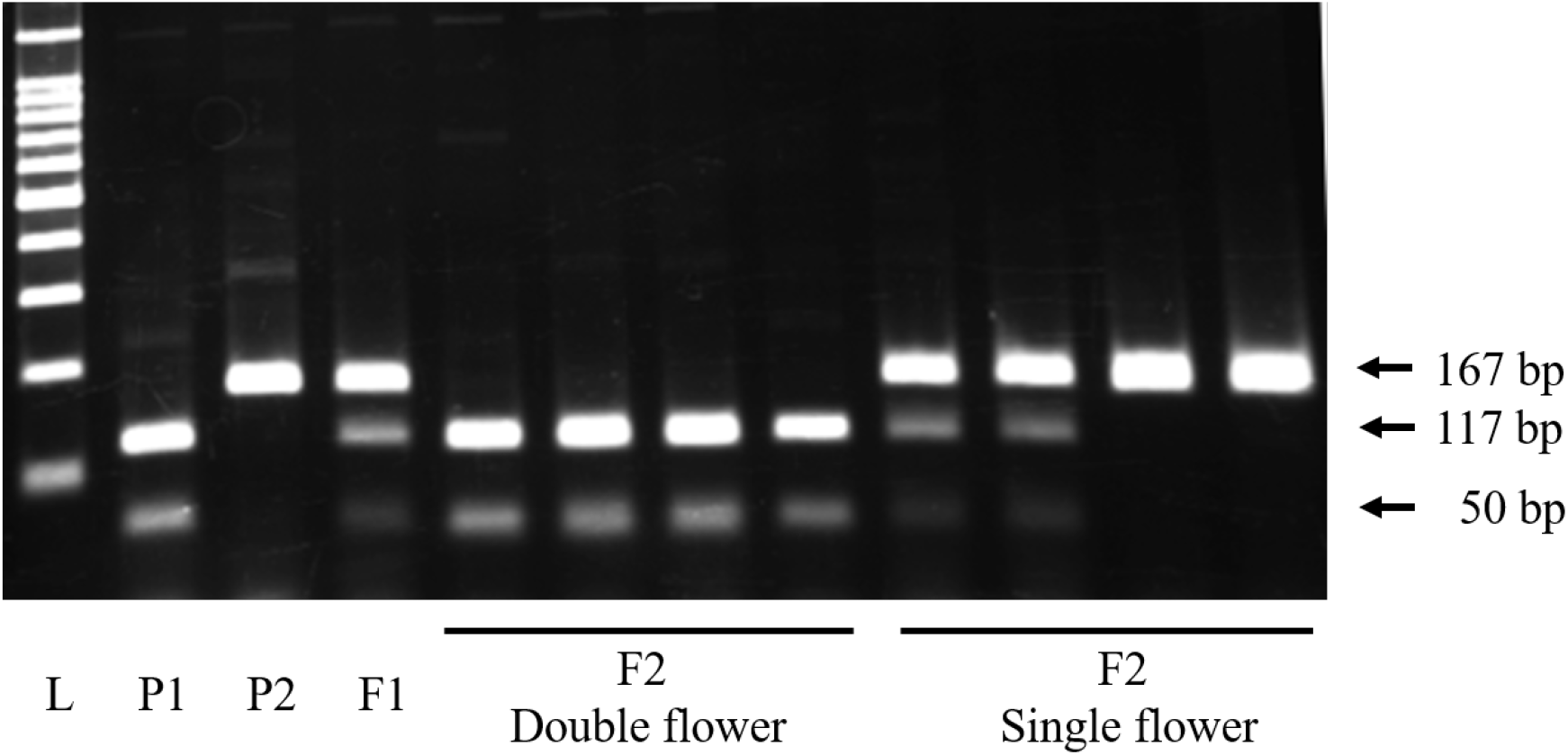
Fragment pattern of J01 DNA marker Dominant single flower allele is shown as undigested 167 bp fragment. Recessive double flower allele is shown as digested 117 and 50 bp fragments. L: 100 bp ladder, P1: ‘Posy Bouquet Grace’ (117_50/117_50), P2: ‘Blue Picotee Manaslu’ (167/167).

**Figure 4.**
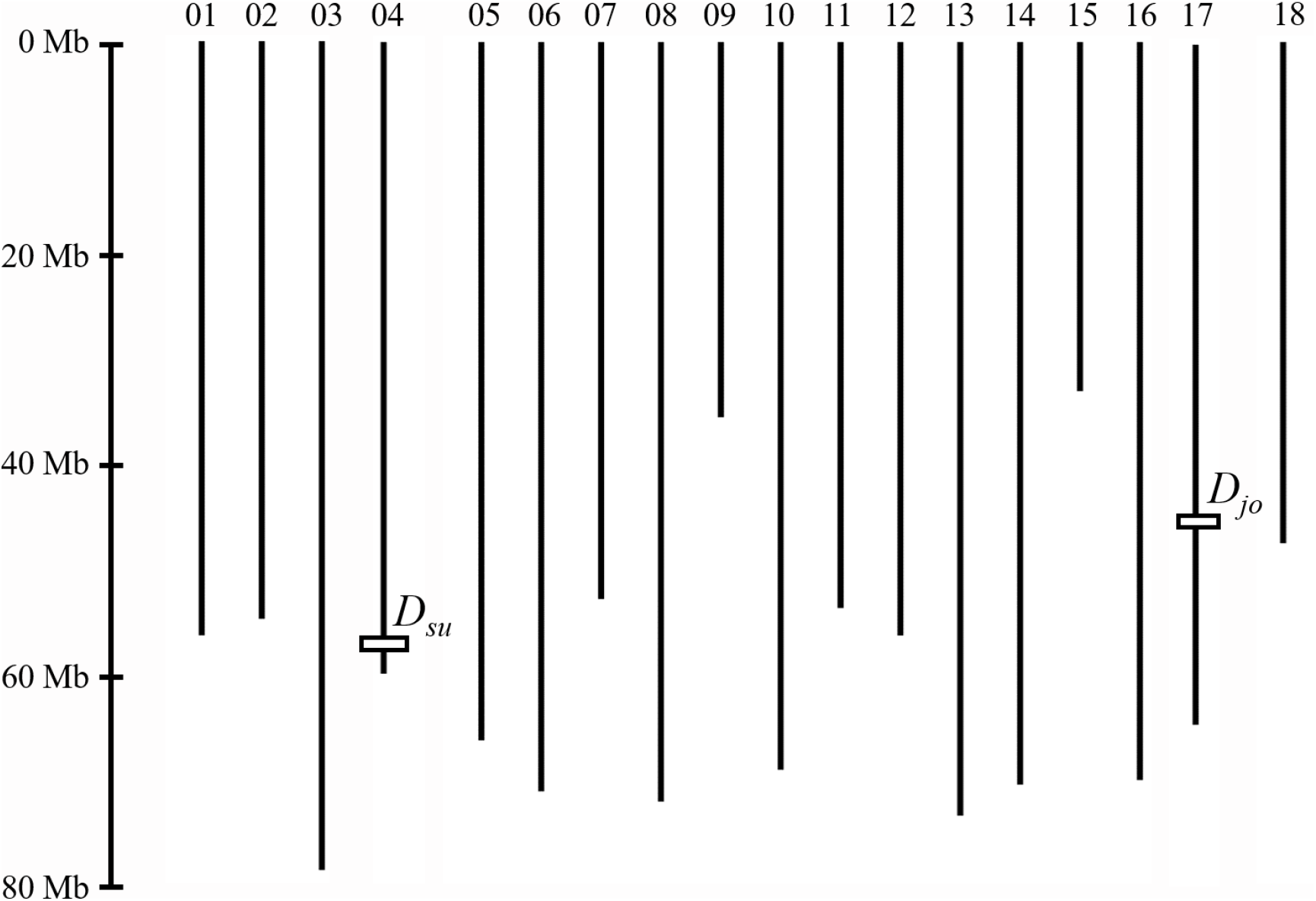
Schematic model of pseudomolecules Double flower phenotype controlling loci *D*_*su*_ and *D*_*jo*_ are shown. *D*_*jo*_ is shown as J01 marker position 46,326,384 in CHR17. *D*_*su*_ is shown as tightly linked SNP at 0109F_868569, since the S01 marker sequence was not on the pseudomolecule.

For identification of SNPs linked to the double flower phenotype of ‘Sumidanohanabi,’ the KF population that segregates the double flower phenotype derived from ‘Sumidanohanabi’ were used. First, we tried to find co-segregated scaffolds with the double flower phenotype by ddRAD-Seq analysis of the KF population. As a result of ddRAD-Seq analysis, 15,102 of SNPs were called. In this population, the double flower phenotype was expected when the plant was homozygous for the ‘Kirakiraboshi’ genotype, and the single flower phenotype was expected when the plant was homozygous for ‘Frau Yoshimi’ or was heterozygous. Each SNP was tested for its fitting rate to this model. As a result, five SNPs on three scaffolds were found to have more than a 95% fitting rate with the model (Table 2). Since SNPs on scaffold 3145F all had the same genotype across the KF population, three loci—on scaffold 0577F, 3145F, 0109F—were detected. According to genotypes of the KF population, these three loci were tightly linked within 5 cM; 0109F (0 cM) - 3145F (3.9 cM) - 0577F (5.0 cM). Since the SNP at position 868569 in 0109F was found at the position 57,436,162 in CHR04, locus *D*_*su*_, which controls the double flower phenotype of ‘Sumidanohanabi,’ was suggested to be located on terminal of CHR04 (Figure 4).

**Table 2.** SNPs correlated (fitting rate more than 95%) with double flower phenotype in KF population

| Scaffold | Position at Phase 0 | Sequence variant |  | Fitting rate (%) | Frequency of double flower phenotype (double flower/all) |  |  |
| --- | --- | --- | --- | --- | --- | --- | --- |
|  |  | Kirakiraboshi | Frau Yoshimi |  | Homozygous of 'Kirakiraboshi' | Heterozygous | Homozygous of 'Frau Yoshimi' |
| 0577F | 1204837 | AG | AAACATG | 98.9 | 22/22 | 0/51 | 1/20 |
| 3145F | 55089 | TA | TAA | 98.9 | 22/22 | 0/51 | 1/20 |
| 3145F | 55109 | G | A | 98.9 | 22/22 | 0/51 | 1/20 |
| 3145F | 55446 | G | A | 98.9 | 22/22 | 0/51 | 1/20 |
| 0109F | 868569 | C | G | 95.7 | 22/25 | 0/44 | 1/24 |

### 3.5 Prediction of genes controlling double flower

To find the gene controlling *D*_*su*_ and *D*_*jo*_, we searched the homeotic genes on scaffolds shown in Table 1 and Table 2. We did not find any notable homeotic gene controlling flower phenotype for *D*_*jo*_. For *D*_*su*_, the g182220 gene, which encoded a homeotic gene *LFY*, was found on scaffold 0577F. To investigate the possibility that it was the causative gene for *D*_*su*_, sequence variants on *LFY* genomic sequence were searched to identify ‘Kirakiraboshi’ specific mutation, using resequencing data of ‘Kirakiraboshi,’ ‘Frau Yoshimi,’ ‘Posy Bouquet Grace,’ and ‘Blue Picotee Manaslu.’ As a result, five INDELs and six sequence variants were found as ‘Kirakiraboshi’ specific mutations (Figure 5).

**Figure 5.**
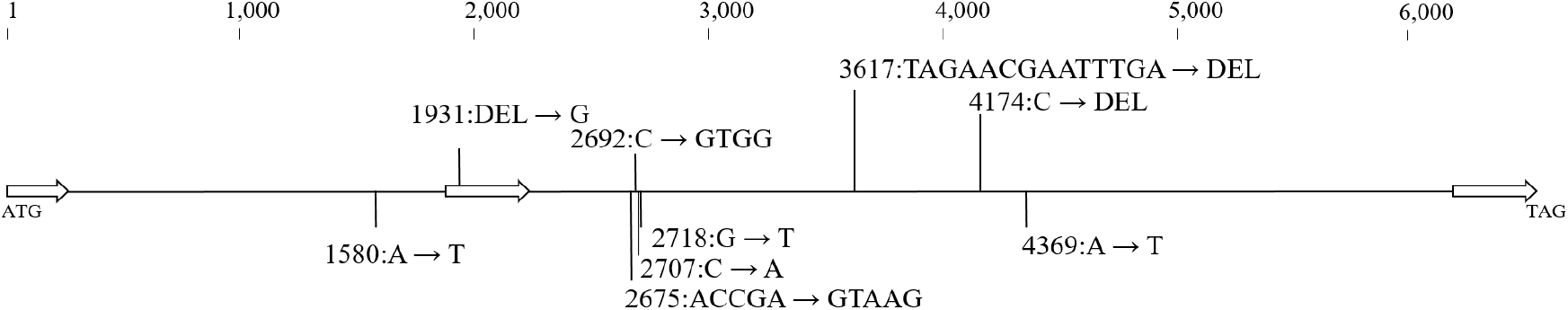
DNA polymorphisms in *LFY* genomic sequence *LFY* sequence polymorphisms observed specifically in ‘Kirakiraboshi’ genomic sequence The sequence is started from the initiation codon (ATG) at 678,200 to the termination signal (TAG) at 684,639 in phase 1 sequence of 0577F of HMA_r1.2. White arrows indicate coding sequences, CDS1: 1 to 454 bp, CDS2: 1,888 to 2,255 bp, CDS3: 6,078 to 6,440 bp. Genetic variants are shown as from Hma1.2 sequence to ‘Kirakiraboshi’.

Cloning and sequencing of *LFY* CDS was performed on ‘Kirakiraboshi’ and ‘Frau Yoshimi.’ From ‘Frau Yoshimi,’ a single CDS comprising three exons was obtained. From ‘Kirakiraboshi,’ two CDSs with splice variants were obtained. While splicing 1 CDS resulted in three exons, splicing 2 CDS resulted in only two exons, corresponding to the first and third splice products of splicing 1 CDS (Supplementary Figure S1). The deduced amino acid sequences were aligned using CDSs of ‘Frau Yoshimi’ and ‘Kirakiraboshi,’ g182220 sequence, protein LFY of *Arabidopsis thaliana*, and protein FLO of *Antirrhium majos*. While the deduced amino acid sequences of ‘Frau Yoshimi’ and g182220 showed sequence similarity in the entire region, frameshift occurred in the two CDSs obtained from ‘Kirakiraboshi’ and the resulting products had no sequence similarity across the latter half (Figure 6). Frameshift observed in splicing 1 CDS was due to one bp of DNA insertion in the second exon, at position 1,931 (Figure N3A). On the contrary, frameshift observed in splicing 2 CDS was due to the complete loss of the second exon (Figure 6).

**Figure 6.**
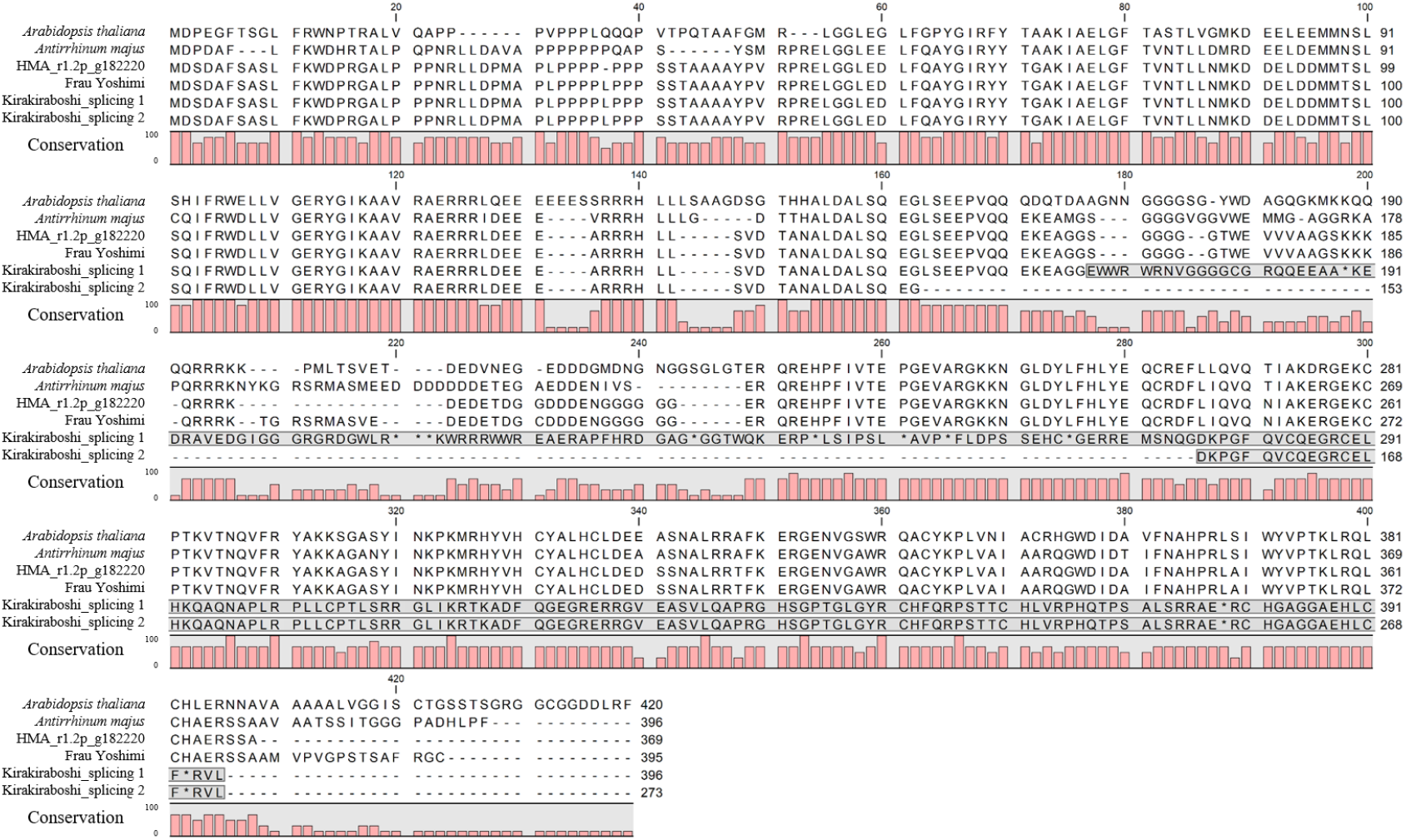
Alignment of LFY protein sequences Amino acids with gray background show frameshifted regions. Splicing variant was observed, and both sequences showed frameshift in ‘Kirakiraboshi’. *Arabidopsis thaliana:* ABE66271.1 *Antirrhium majus*: AAA62574.1.

To develop a DNA marker for distinguishing the *d*_*su*_ allele from the *D*_*su*_ alleles in the *LFY* genomic sequence, we focused and designed a DNA marker on ‘Kirakiraboshi’ specific 14 bp deletion at position 3,617 from initiation codon (Figure 5). We developed INDEL S01 marker amplified 236 bp fragment for the double flower allele of ‘Kirakiraboshi,’ and 250 bp and 280 bp fragments for the single flower allele of ‘Frau Yoshimi’ (Figure 7A). Three types of alleles resulted from the presence or absence of a 30 bp deletion at position 3,482 in addition to the 14 bp INDEL. These were both 30 bp and 14 bp deletions on the 236 bp allele, 30 bp deletion on the 250 bp allele, and no deletion on the 280 bp allele (Figure 7B).

**Figure 7.**
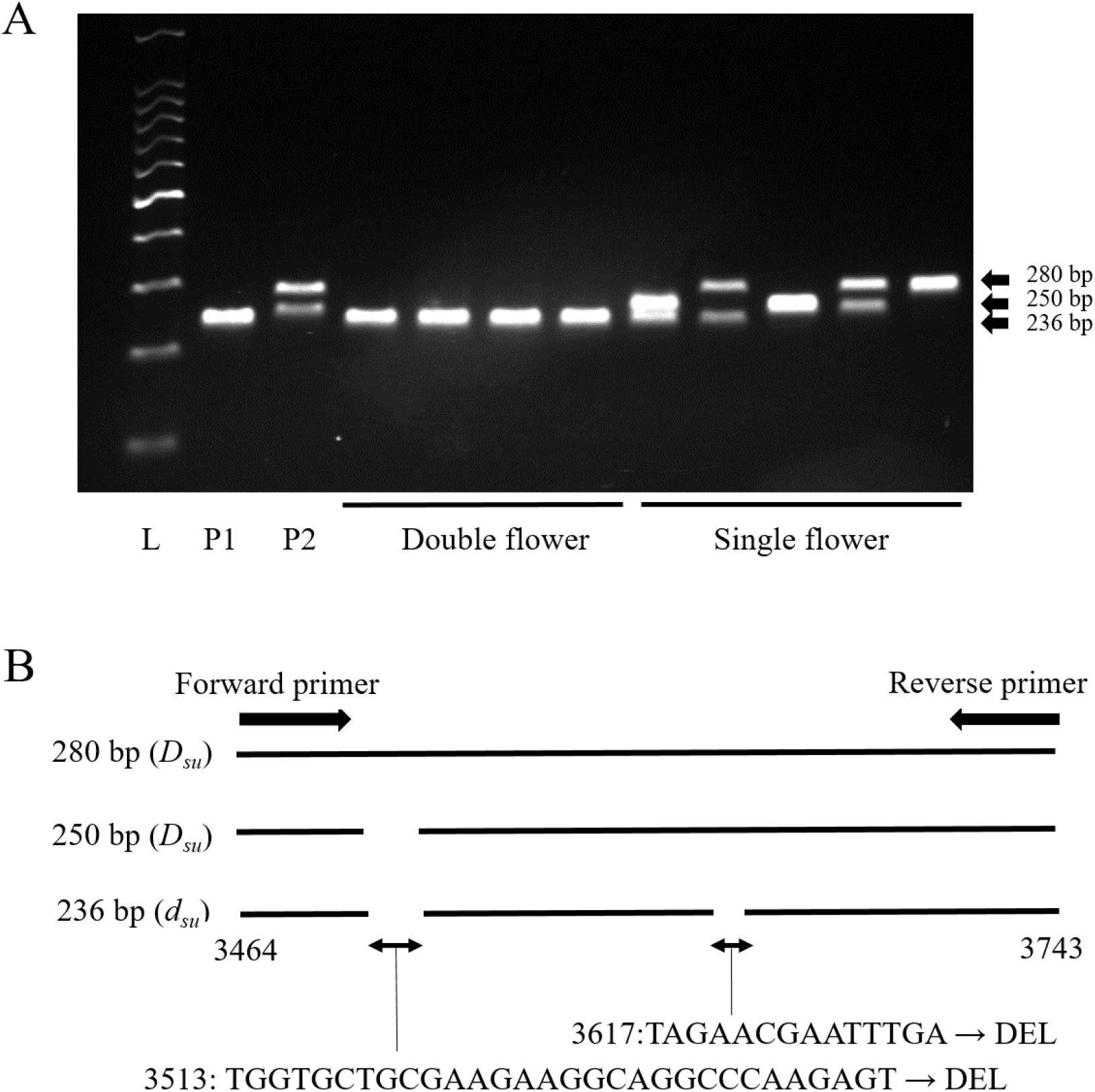
Fragment pattern of S01 DNA marker Fragment pattern of S01 DNA marker. Dominant single flower alleles are shown as 250 bp and 280 bp fragments. Recessive double flower allele is shown as 236 bp fragments. L: 100 bp ladder, P1: ‘Kirakiraboshi’ (236/236), P2: ‘Frau Yoshimi’ (250/280). INDEL polymorphisms in alleles of DNA marker S01 amplified sequences. Position on schematic models were the same as in Figure 5.

### 3.6 Genotyping of hydrangea accessions using J01 and S01 markers

Since the J01 marker could distinguish *D*_*jo*_/*d*_*jo*_ alleles and the S01 marker could distinguish *D*_*su*_/*d*_*su*_ alleles, a combined use of J01 and S01 DNA markers was expected to reveal the origin of the double flower phenotype, *d*_*jo*_ or *d*_*su*_, in various accessions. Therefore, DNA marker genotyping on *H. macrophylla* accessions were performed using two DNA markers, J01 and S01. All tested double flower accessions showed homozygous genotypes of J01 or S01; ten of the double flower accessions were homozygous of 117_50 in J01, and four were homozygous of 236 in S01 (Table 3). Contrarily, all single flower accessions showed other genotypes.

**Table 3.** Genotypes of DNA marker J01 and S01 in *H. macrophylla* varieties

| Accession name | Phenotype | Genotype |  |
| --- | --- | --- | --- |
|  |  | J01 | S01 |
| Jogasaki | Double | 117_50/117_50 | 250/280 |
| Posy Bouquet Grace | Double | 117_50/117_50 | 280/280 |
| Izunohana | Double | 117_50/117_50 | 250/280 |
| Chikushinokaze | Double | 117_50/117_50 | 250/280 |
| Chikushinomai | Double | 117_50/117_50 | 280/280 |
| Chikushiruby | Double | 117_50/117_50 | 280/280 |
| Corsage | Double | 117_50/117_50 | 280/280 |
| Dance Party | Double | 117_50/117_50 | 280/280 |
| Fairy Eye | Double | 117_50/117_50 | 250/280 |
| Posy Bouquet Casey | Double | 117_50/117_50 | 250/280 |
| Sumidanohanabi | Double | 167/167 | 236/236 |
| Kirakiraboshi | Double | 167/167 | 236/236 |
| HK01 | Double | 167/167 | 236/236 |
| HK02 | Double | 167/167 | 236/236 |
| 03JP1 | Single | 117_50/167 | 280/280 |
| Amethyst | Single | 167/167 | 250/280 |
| Blue Picotee Manaslu | Single | 167/167 | 280/280 |
| Blue Sky | Single | 167/167 | 280/280 |
| Bodensee | Single | 167/167 | 250/250 |
| Chibori | Single | 167/167 | 280/280 |
| Furau Mariko | Single | 167/167 | 250/250 |
| Furau Yoshiko | Single | 167/167 | 280/280 |
| Furau Yoshimi | Single | 167/167 | 250/280 |
| Green Shadow | Single | 167/167 | 280/280 |
| Kanuma Blue | Single | 167/167 | 250/280 |
| Mrs. Kumiko | Single | 167/167 | 280/280 |
| Paris | Single | 167/167 | 280/280 |
| Peach Hime | Single | 167/167 | 280/280 |
| Picotee | Single | 167/167 | 282/282 |
| Ruby Red | Single | 167/167 | 280/280 |
| Shinkai | Single | 167/167 | 280/280 |
| Tokimeki | Single | 167/167 | 280/282 |
| Uzuajisai | Single | 167/167 | 250/280 |

Previously, the double flower phenotype has been revealed to be controlled by a single locus with the inheritance of single flower dominant and double flower recessive genes^4,5^. It was also suggested that genes controlling the double flower phenotype were different between ‘Jogasaki’ and ‘Sumidanohanabi’ based on confirmation of the segregation ratio of crossed progenies^4^. Our study revealed that the double flower phenotype of ‘Jogasaki’ was controlled by a single *D*_*jo*_ locus on CHR17, and the double flower phenotype of ‘Sumidanohanabi’ was controlled by a single *D*_*su*_ locus on CHR04. In addition, all double flower accessions showed homozygosity for the double flower allele at one locus, *D*_*jo*_ or *D*_*su*_. Contrarily, all single flowers have dominant single flower alleles on both *D*_*jo*_ and *D*_*su*_ loci. This indicated that each locus independently controls flower phenotype.

Developed DNA markers J01 and S01 could successfully identify recessive double flower alleles for *D*_*jo*_ and *D*_*su*_, respectively. Both markers showed high fitting ratio with phenotype and were applicable to the examined *H. macrophylla* accessions. The S01 marker is superior to the DNA marker STAB045 linked to *D*_*su*_ and which was discovered by Waki et al.^5^ because the former has a wide range of applicability. While the S01 marker genotype completely fitted with the phenotype in all tested accessions, STAB045 did not (data not shown). Because both J01 and S01 showed a wide range of applicability, it is advantageous to use them in combination to reveal the existence of the double flower allele in *H. macrophylla* accessions. This information will help in selection of candidate parents with heterozygous recessive double flower alleles to obtain double flower progenies. In addition, these DNA markers should be useful in marker assisted selection (MAS) of double flower progenies. To obtain double flower progenies, at least the paternal parent should be of the single flower phenotype because very few or none at all pollen grains are produced in double flower individuals. In addition, it requires approximately 2 years to confirm the flower phenotype from the time of crossing. Identification of flower phenotype at the seedling stage by MAS would enable the discarding of single flower individuals and allow the growth of double flower individuals. The developed DNA markers should accelerate the breeding of double flower phenotypes.

In the genomic sequence of ‘Kirakiraboshi,’ an insertion was detected in the second exon of the *LFY* gene. This insertion actually resulted in frameshift of cloned mRNA in both splice variants. Therefore, it was speculated that the function of the *LFY* gene was suppressed or lost in ‘Kirakiraboshi’. The *LFY* gene and its homologue *FLO* have been identified in many plants, such as *Arabidopsis thaliana* and *Antirrhinum majus*, and are known as transcription factors for major flowering signals^29–31^. Additionally, many types of phenotypes in *Arabidopsis lfy* mutants have been reported^32,33^. In the *lfy* strong phenotype, most organs are sepal-like, or mosaic sepal/carpels organs, and the sepal-like organs are characteristic of wild-type cauline leaves^33^. Therefore, the flowers of the *lfy* mutant appeared to be double flowers that are formed from leaves or sepals. Additionally, a similar phenotype has been reported in *LFY* homologue mutants or transgenic plants such as the *flo* mutant of *Antirrhinum majus*^34^, *uni* mutant of pea^35^, and co-suppressed *NFL* transgenic plant of tobacco^37^. Therefore, generally, when the *LFY* gene function is lost, petal, stamens, and a carpel are likely to be replaced by sepal-like organs. In decorative flowers of hydrangea, sepals show petaloid characteristics including pigmentation and enlarged organ size. It is possible that sepal-like organs in decorative flowers show petaloid characteristics and form double flowers. Therefore, we assumed that *LFY* is a causative gene of the double flower phenotype of ‘Sumidanohanabi’.

However, there remain several unexplained observations in this study. The double flower of ‘Kirakiraboshi’ did not exhibit the exact same phenotype of the *lfy* mutant. Generally, the flowers of *lfy* or its orthologous gene mutants have only leaf-like or sepal-like organs that have chlorophyll, stomata, and trichome, and these organs have almost no petal identity^33,34^. When flowering signals in *lfy* mutant were lost completely, floral organs were not fully formed^33-35^. In the double flowers of ‘Kirakiraboshi’, the floral organs keep their petal identity, have papilla cells, and are pink or blue.

These phenotypes of ‘Kirakiraboshi’ might reflect partial remaining of LFY function. Additionally, it has been reported that *lfy* mutants with an intermediate or weak phenotype sometimes develop petaloid organs^33^. According to the genomic sequence of *H. macrophylla*, no other *LFY* gene was observed. It could be considered that the double flowers of ‘Kirakiraboshi’ were induced via partial repression of the LFY function.

On the contrary, we could not find any candidate gene that controls the double flower phenotype for the *D*_*jo*_ locus. One possible reason was that SNPs were not called in scaffold with causative gene. In pseudomolecules, about half of the total scaffolds length was not included since relevant SNPs were not called. Improvement of SNP density would be effective for discovering additional scaffolds that are tightly linked to *D*_*jo*_. Although candidate gene for *D*_*jo*_ could not be identified from the linkage information, we predicted several candidate genes. In hydrangea, stamens and petals were absent from decorative flowers of the double flower plant, and there was an increased number of sepals^4^. Since causative genes should explain the changes in formation, the B-class genes of the ABC model, *PI* and *AP3*, were predicted as candidate genes. In *A. thaliana*, the B-class gene *pi* or *ap3* mutants showed an increase in the number of sepals converted from petals^37^. If these genes were mutated in hydrangea, an increase in sepals would be expected. In hydrangea, *HmPI*, *HmAP3*, and *HmTM6* were identified as B-class genes^38,39^. As *HmAP3* was located on CHR13, it was not considered as a causative gene for *D_jo_*. In this study, *HmPI* and *HmTM6* were not included in the pseudomolecule. Ascertaining the loci of these genes might reveal the causative gene for *D*_*jo*_.

In this study, we report DNA markers and possible causative genes for the double flower phenotype observed in two hydrangea cultivars. For this analysis, we established a reference sequence for the hydrangea genome using advanced sequencing technologies including the long-read technology (PacBio) and the HiC method^9^, bioinformatics techniques for the diploid genome assembly^14^, and haplotype phasing^8^. To the best of our knowledge, this is the first report on the chromosome-level haplotype-phased sequences in hydrangea at the level of the species (*H. macrophylla*), genus (*Hydrangea*), family (Hydrangeaceae), and order (Cornales). The genomic information from this study based on NGS technology is a significant contribution to the genetics and breeding of hydrangea and its relatives. It will serve to accelerate the knowledge base of the evolution of floral characteristics in Hydrangeaceae.

## Acknowledgments

We thank Ohama A, Ono M, Seki A and Kitagawa A (Nihon University) and Sasamoto S, Watanabe A, Nakayama S, Fujishiro T, Kishida Y, Kohara M, Tsuruoka H, Minami C, and Yamada M (Kazusa DNA Research Institute) for their technical help.

## Funding

This study was partially supported by the Nihon University College of Bioresource Sciences Research Grant for 2018, and by the JSPS KAKENHI Grant, Number JP18K14461.

## Supporting information

**Supplementary Table S1.** RNA samples used for Iso-Seq and RNA-Seq

**Supplementary Table S2.** Statistics of the genome sequences of *Hydrangea macrophylla* ‘Aogashima-1’

**Supplementary Table S3.** J01 marker genotypes and double flower phenotypes of 15IJP1 population.

**Supplementary Table S4.** J01 marker genotypes and double flower phenotypes of 14GT77 population.

**Supplementary Figure S1.** Alignment of *LFY* genomic sequence and CDS.

## Data availability

The sequence reads are available from the DNA Data Bank of Japan (DDBJ) Sequence Read Archive (DRA) under the accession numbers DRA010300, DRA010301, and DRA010302. The assembled sequences are available from the BioProject accession number PRJDB10054. The genome information is available at Plant GARDEN (https://plantgarden.jp).

## Notes

### Competing Interest Statement

The authors have declared no competing interest.

## References

1. Uemachi, T., Kato, Y., and Nishio, T. 2004, Comparison of decorative and non-decorative flowers in *Hydrangea macrophylla* (Thunb.) Ser., Sci. Hortic., 102, 325–334

2. Uemachi, T., Kurokawa, M., and Nishio, T. 2006, Comparison of inflorescence composition and development in the lacecap and its sport, hortensia *Hydrangea macrophylla* (Thunb.) Ser., J. Japan. Soc. Hort. Sci., 75, 154–160.

3. Uemachi, T. and Okumura, A. 2012, The inheritance of inflorescence types in *Hydrangea macrophylla*, J. Japan. Soc. Hort. Sci., 81, 263–268.

4. Suyama, T., Tanigawa, T., Yamada, A. et al. 2015, Inheritance of the double-flowered trait in decorative hydrangea flowers, Hortic. J., 84, 253–260.

5. Waki, T., Kodama, M., Akutsu, M. et al. 2018, Development of DNA markers linked to double-flower and hortensia traits in *Hydrangea macrophylla* (Thunb.) Ser., Hortic J., 87, 264–273.

6. Heijmans, K., Ament, K., Rijpkema, A.S. et al. 2012, Redefining C and D in the petunia ABC, Plant Cell, 24, 2305–2317.

7. Tränkner, C., Krüger, J., Wanke, S., Naumann, J., Wenke, T. and Engel, F. 2019, Rapid identification of inflorescence type markers by genotyping-by-sequencing of diploid and triploid F1 plants of *Hydrangea macrophylla*, BMC Genet., 20, 60.

8. Kronenberg, Z. N., Hall, R. J., Hiendleder, S., et al. 2018, FALCON-Phase: Integrating PacBio and Hi-C data for phased diploid genomes, BioRxiv, 327064.

9. Dudchenko, O., Batra, S. S., Omer, A. D., et al. 2017, De novo assembly of the *Aedes aegypti* genome using Hi-C yields chromosome-length scaffolds. Science, 356, 92–95.

10. Mascher, M. and Stein, N. 2014, Genetic anchoring of whole-genome shotgun assemblies, Front Genet, 5, 208.

11. Marcais, G. and Kingsford, C. 2011, A fast, lock-free approach for efficient parallel counting of occurrences of k-mers, Bioinformatics, 27, 764–770.

12. Kajitani, R., Toshimoto, K., Noguchi, H., et al. 2014, Efficient de novo assembly of highly heterozygous genomes from whole-genome shotgun short reads, Genome Res, 24, 1384–1395.

13. Simao, F. A., Waterhouse, R. M., Ioannidis, P., Kriventseva, E. V., and Zdobnov, E. M. 2015, BUSCO: assessing genome assembly and annotation completeness with single-copy orthologs, Bioinformatics, 31, 3210–3212.

14. Chin, C. S., Peluso, P., Sedlazeck, F. J., et al. 2016, Phased diploid genome assembly with single-molecule real-time sequencing, Nat Methods, 13, 1050–1054.

15. Walker, B. J., Abeel, T., Shea, T., et al. 2014, Pilon: an integrated tool for comprehensive microbial variant detection and genome assembly improvement, PLoS One, 9, e112963.

16. Shirasawa, K., Hirakawa, H., and Isobe, S. 2016, Analytical workflow of double-digest restriction site-associated DNA sequencing based on empirical and in silico optimization in tomato, DNA Res, 23, 145–153.

17. Rastas, P. 2017, Lep-MAP3: robust linkage mapping even for low-coverage whole genome sequencing data, Bioinformatics, 33, 3726–3732.

18. Tang, H., Zhang, X., Miao, C., et al. 2015, ALLMAPS: robust scaffold ordering based on multiple maps, Genome Biol, 16, 3.

19. Grabherr, M. G., Haas, B. J., Yassour, M., et al. 2011, Full-length transcriptome assembly from RNA-Seq data without a reference genome, Nat Biotechnol, 29, 644–652.

20. Stanke, M., Keller, O., Gunduz, I., Hayes, A., Waack, S., and Morgenstern, B. 2006, AUGUSTUS: ab initio prediction of alternative transcripts, Nucleic Acids Res, 34, W435–439.

21. Kent, W. J., 2002, BLAT - the BLAST-like alignment tool, Genome Res, 12, 656–664.

22. Ghelfi, A., Shirasawa, K., Hirakawa, H., and Isobe, S. 2019, Hayai-Annotation Plants: an ultra-fast and comprehensive functional gene annotation system in plants, Bioinformatics, 35, 4427–4429.

23. Bolger, A.M., Lohse, M., and Usadel, B. 2014, Trimmomatic: a flexible trimmer for Illumina sequence data, Bioinformatics, 30, 2114–2120.

24. Li, H., Handsaker, B., Wysoker, A., et al. 2009, The Sequence Alignment/Map format and SAMtools, Bioinformatics, 25, 2078–2079.

25. Untergasser, A., Cutcutache, I., Koressaar, T. et al. 2012, Primer3--new capabilities and interfaces. Nucleic Acids Res., 40, e115.

26. Schmieder, R. and Edwards, R. 2011, Quality control and preprocessing of metagenomic datasets, Bioinformatics, 27, 863–864.

27. Langmead, B. and Salzberg, S. L. 2012, Fast gapped-read alignment with Bowtie 2, Nat Methods, 9, 357–359.

28. Danecek, P., Auton, A., Abecasis, G., et al. 2011, The variant call format and VCFtools, Bioinformatics, 27, 2156–2158.

29. Jaeger, K.E., Pullen, N., Lamzin, S., Morris, R.J., and Wigge, P.A. 2013, Interlocking feedback loops govern the dynamic behavior of the floral transition in Arabidopsis, Plant Cell, 25, 820–833.

30. Krizek, B.A. and Fletcher, J.C. 2005, Molecular mechanisms of flower development: an armchair guide, Nat. Rev. Genet., 6, 688.

31. William, D.A., Su, Y., Smith, M.R., Lu, M., Baldwin, D.A., and Wagner, D. 2004, Genomic identification of direct target genes of LEAFY, Proc. Nat. Acad. Sci., 101, 1775–1780.

32. Okamuro, J.K., Den Boer, B.G., and Jofuku, K.D. 1993, Regulation of Arabidopsis flower development, Plant Cell, 5, 1183–1193.

33. Weigel, D., Alvarez, J., Smyth, D.R., Yanofsky, M.F., and Meyerowitz, E.M. 1992, LEAFY controls floral meristem identity in Arabidopsis, Cell, 69, 843–859.

34. Carpenter, R. and Coen, E.S. 1990, Floral homeotic mutations produced by transposon-mutagenesis in Antirrhinum majus, Gene. Dev., 4, 1483–1493.

35. Hofer, J., Turner, L., Hellens, R. et al. 1997, UNIFOLIATA regulates leaf and flower morphogenesis in pea, Curr. Biol., 7, 581–587.

36. Ahearn, K.P., Johnson, H.A., Weigel, D., and Wagner, D.R. 2001, NFL1, a *Nicotiana tabacum* LEAFY-like gene, controls meristem initiation and floral structure, Plant Cell Physiol., 42, 1130–1139.

37. Bowman, J.L., Smyth, D.R., and Meyerowitz, E.M. 1989, Genes directing flower development in Arabidopsis, Plant Cell, 1, 37–52.

38. Kitamura, Y., Hosokawa, M., Uemachi, T., and Yazawa, S. 2009, Selection of ABC genes for candidate genes of morphological changes in hydrangea floral organs induced by phytoplasma infection, Sci. Hort., 122, 603–609.

39. Kramer, E.M. and Irish, V.F. 2000. Evolution of the petal and stamen development programs: Evidence from comparative studies of the lower eudicots and basal angiosperms, Int. J. Plant Sci., 161, s29–s40

