## Supplementary Figure S1 for "Genome sequence of *Hydrangea macrophylla* and its application in analysis of the double flower phenotype"

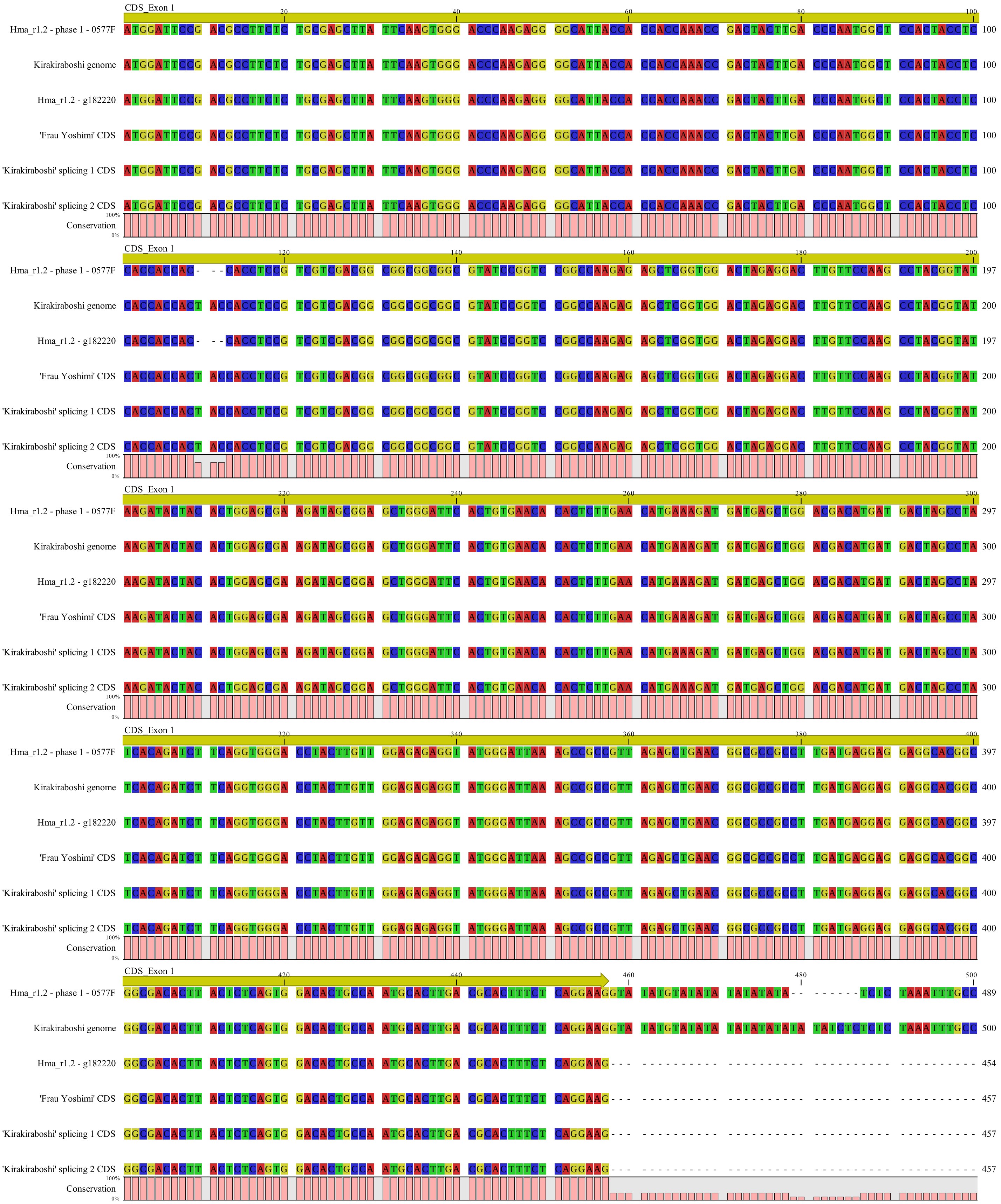


Supplementary Figure S1. Alignment of *LFY* genomic sequence and CDS.

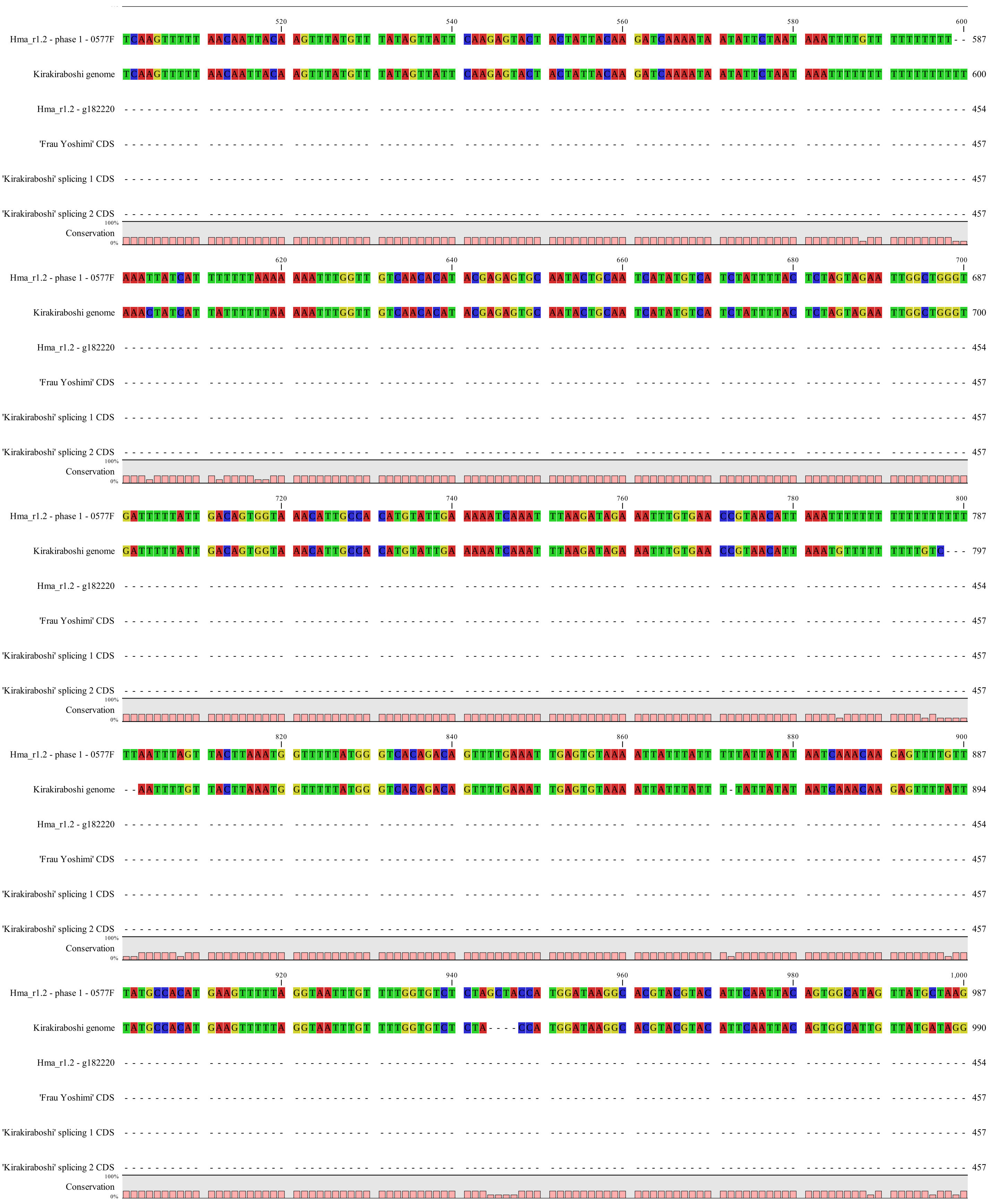


Supplementary Figure S1. Alignment of *LFY* genomic sequence and CDS. (continued)

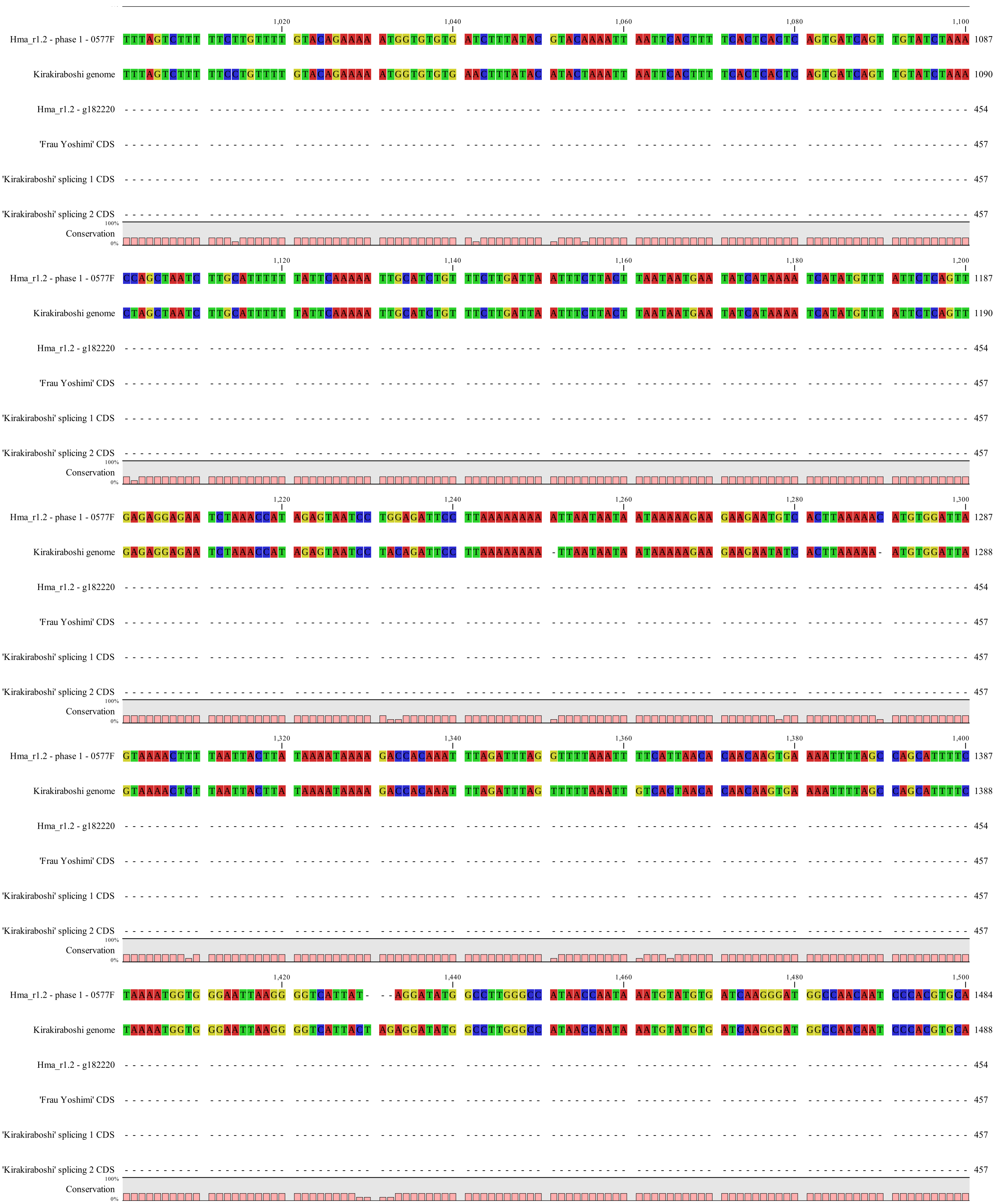

Supplementary Figure S1. Alignment of *LFY* genomic sequence and CDS. (continued)

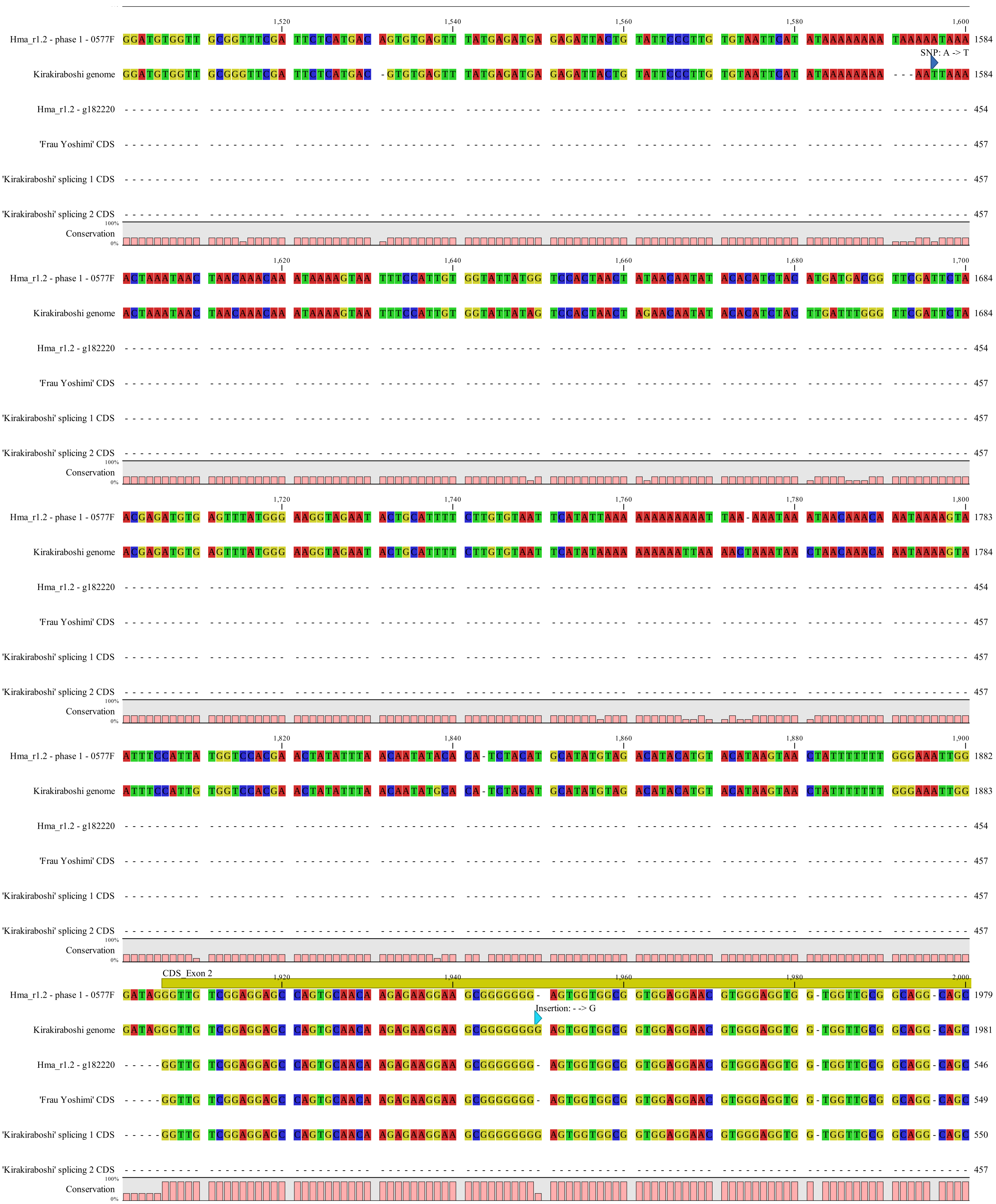


Supplementary Figure S1. Alignment of *LFY* genomic sequence and CDS. (continued)

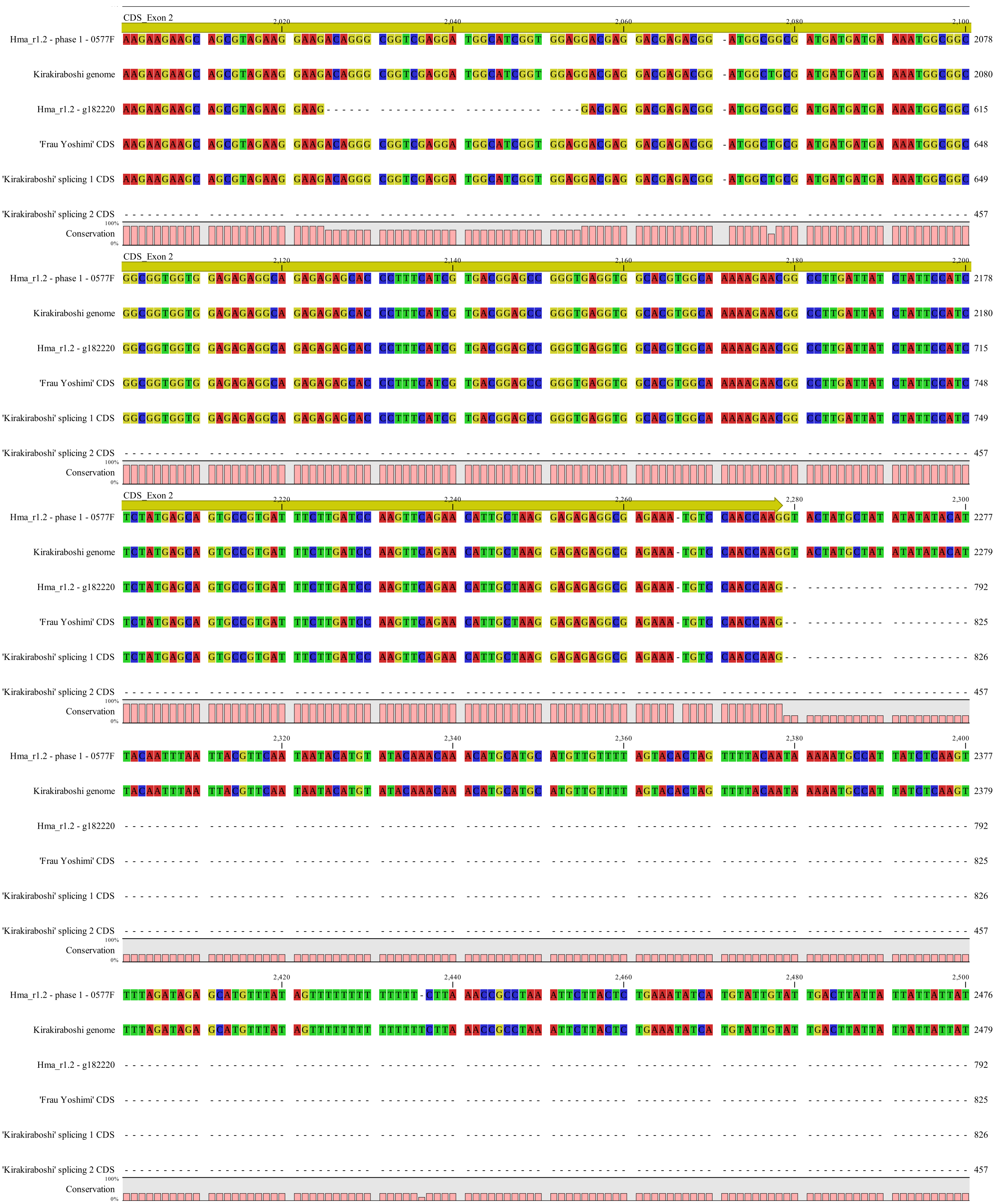


Supplementary Figure S1. Alignment of *LFY* genomic sequence and CDS. (continued)

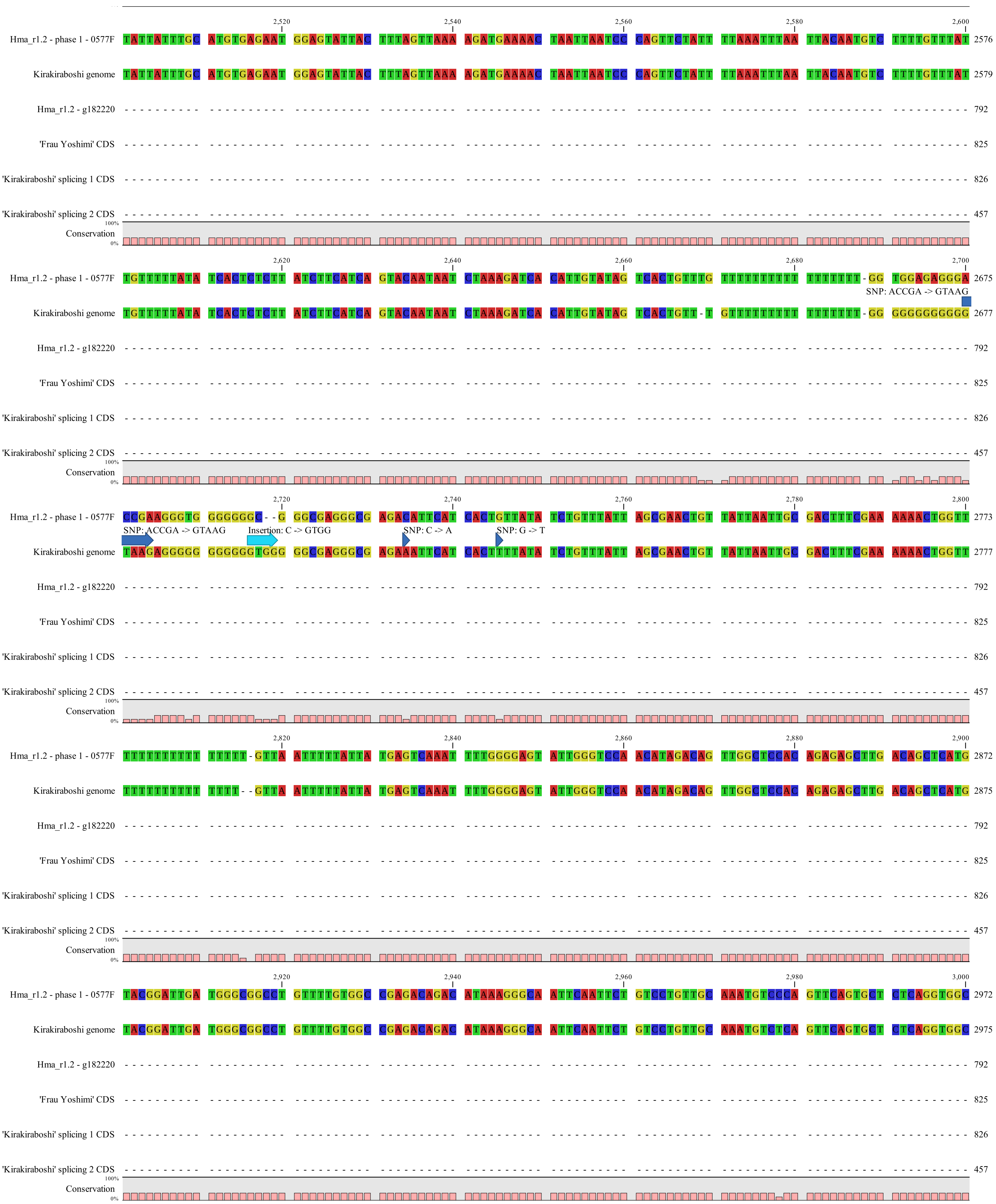


Supplementary Figure S1. Alignment of *LFY* genomic sequence and CDS. (continued)

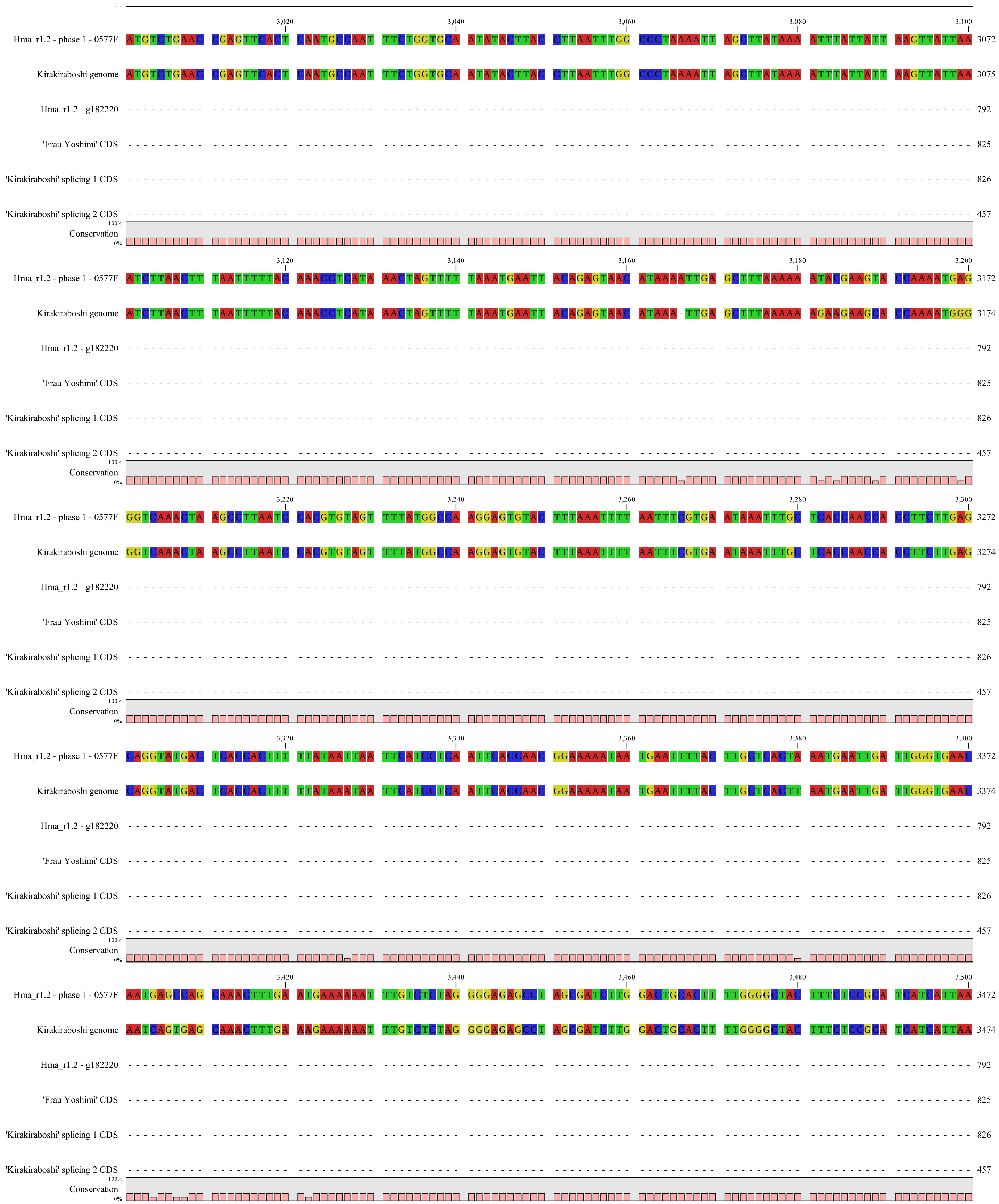


Supplementary Figure S1. Alignment of *LFY* genomic sequence and CDS. (continued)

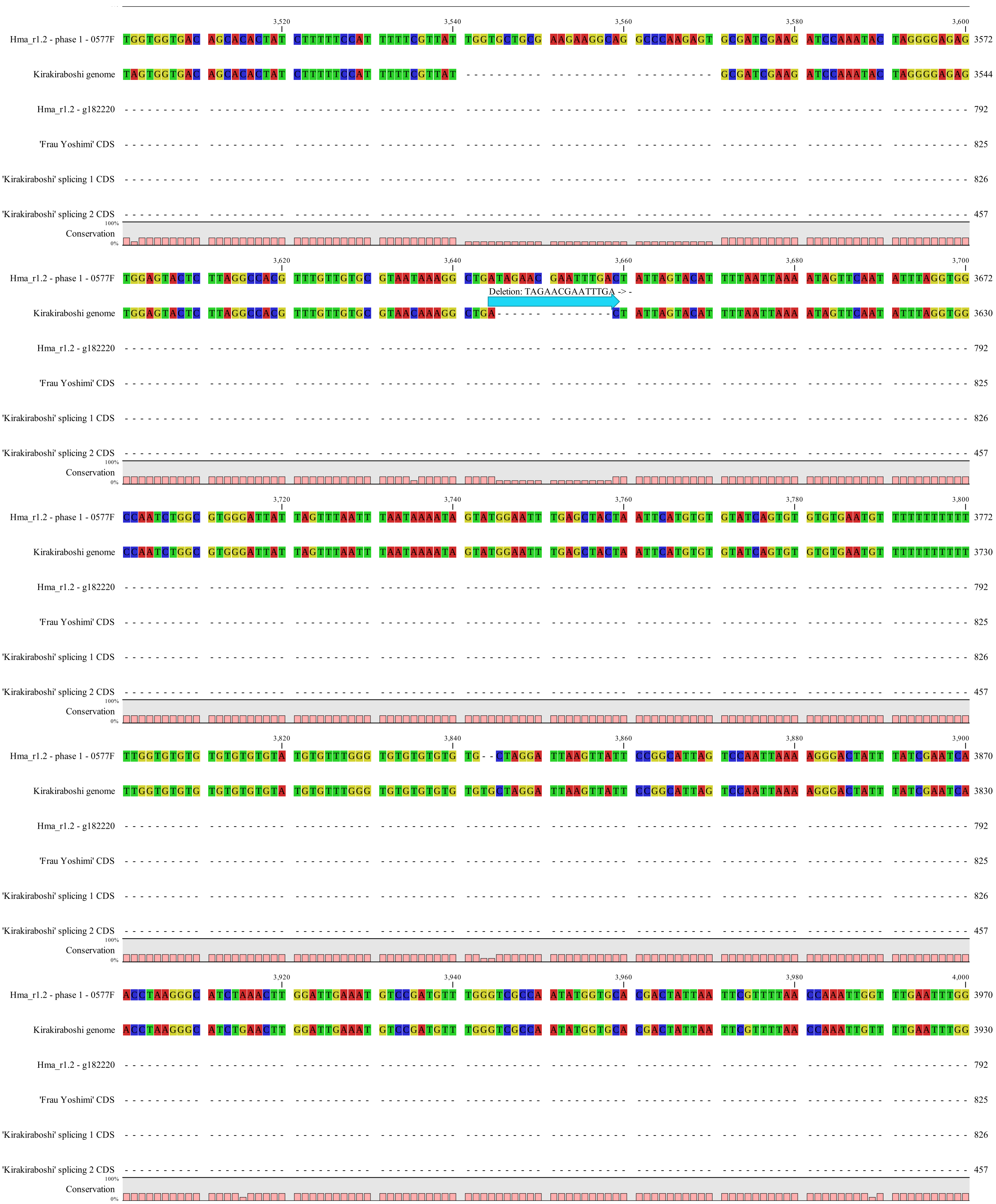


Supplementary Figure S1. Alignment of *LFY* genomic sequence and CDS. (continued)

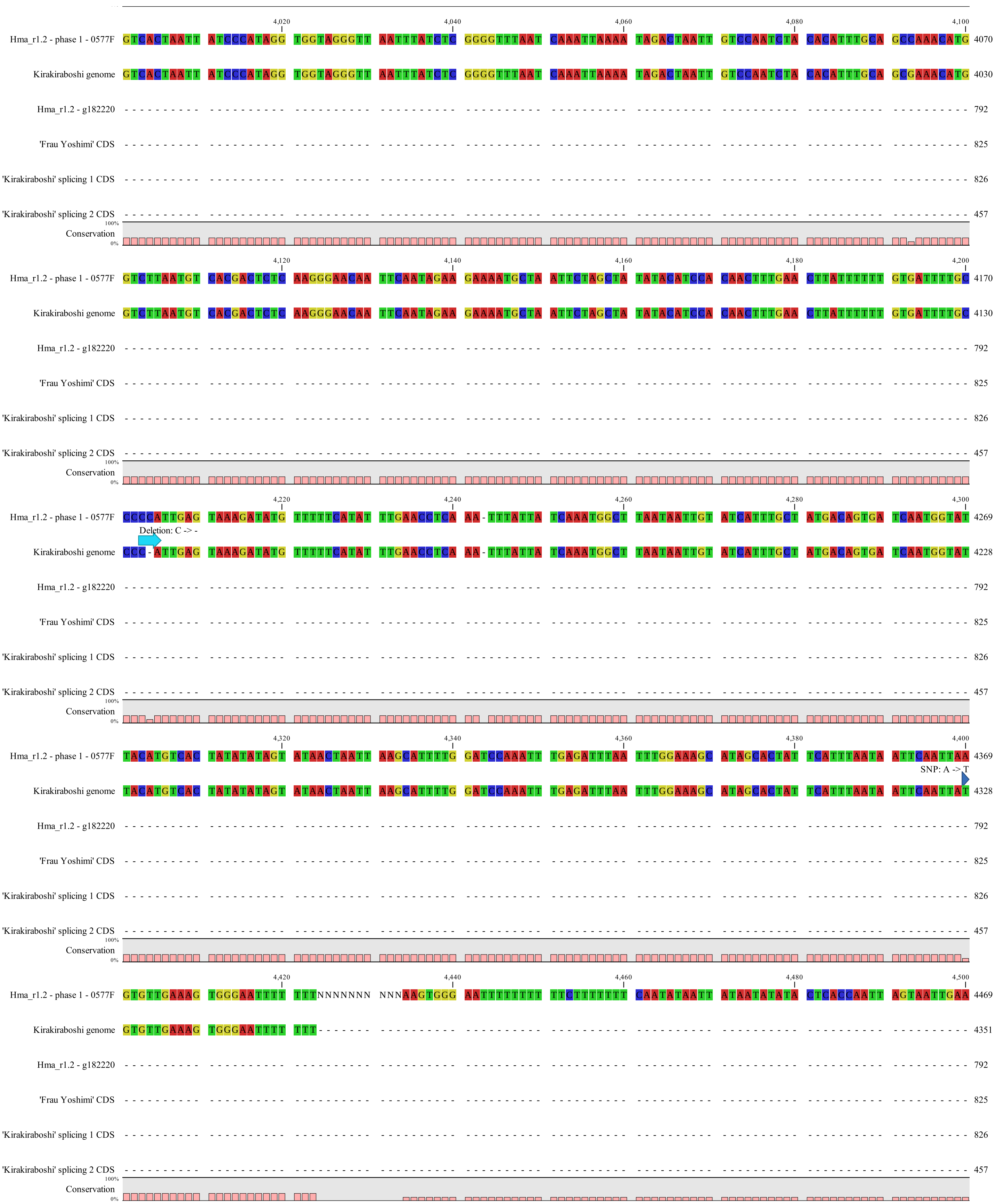


Supplementary Figure S1. Alignment of *LFY* genomic sequence and CDS. (continued)

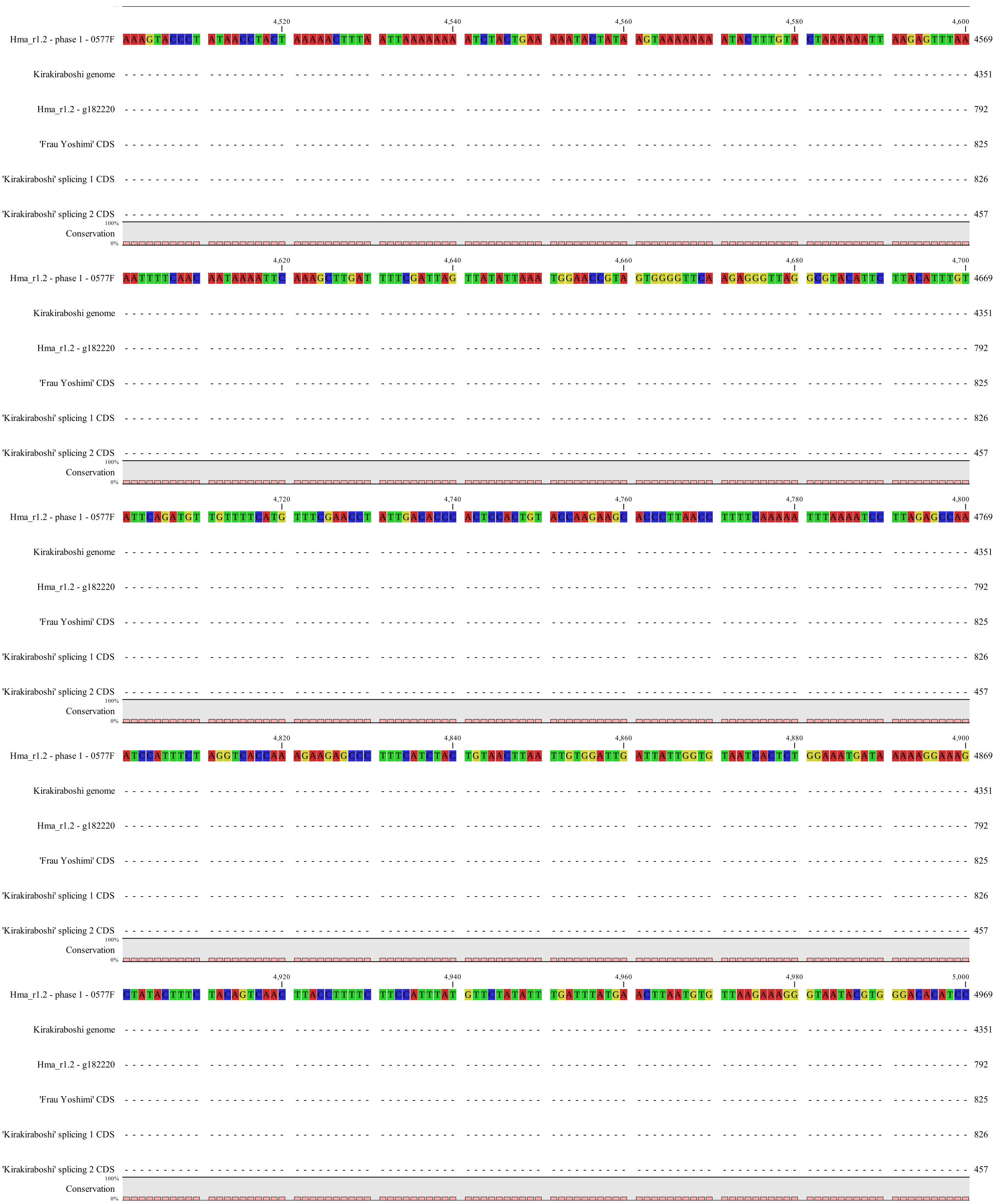


Supplementary Figure S1. Alignment of *LFY* genomic sequence and CDS. (continued)

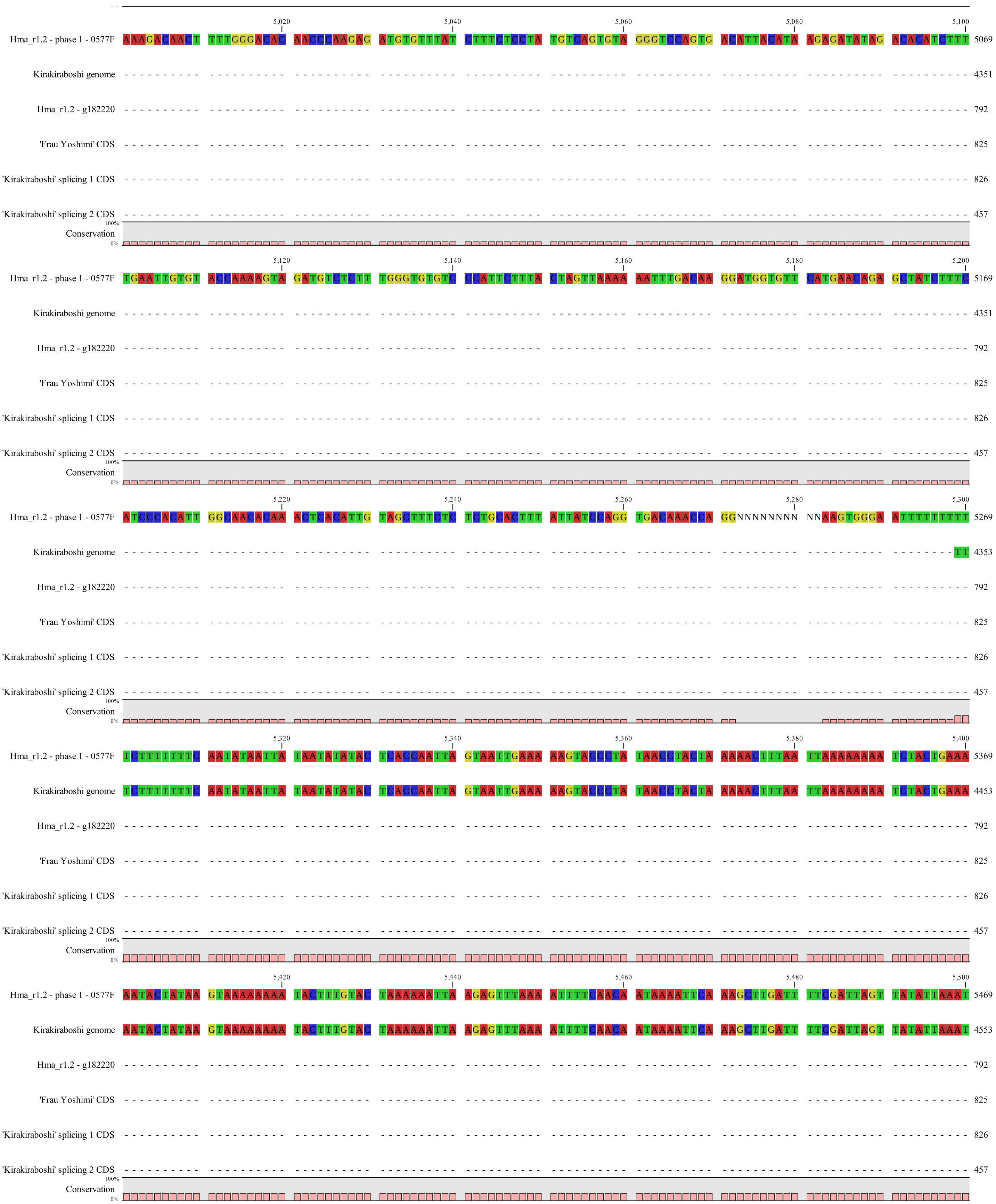


Supplementary Figure S1. Alignment of *LFY* genomic sequence and CDS. (continued)

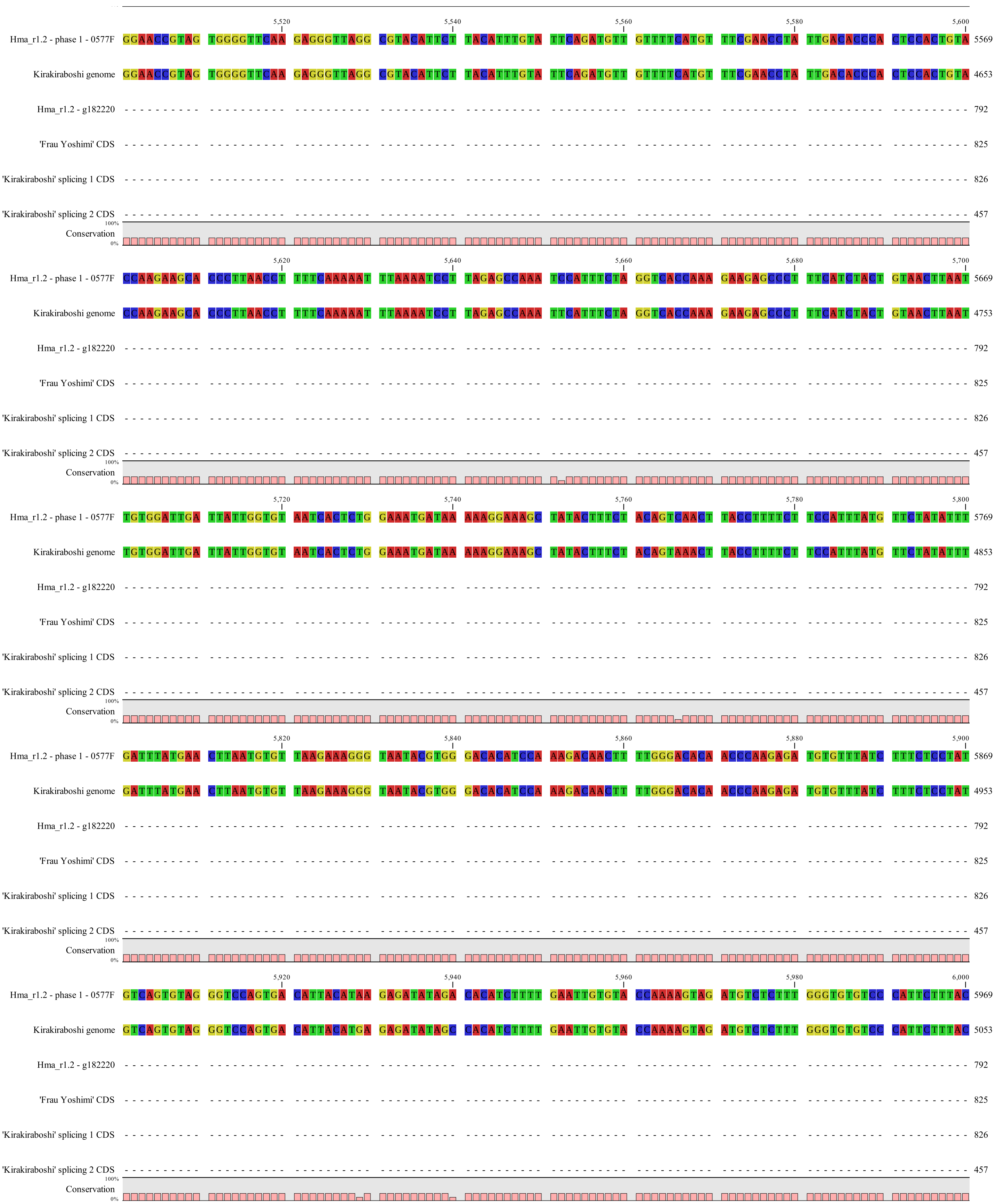


Supplementary Figure S1. Alignment of *LFY* genomic sequence and CDS. (continued)

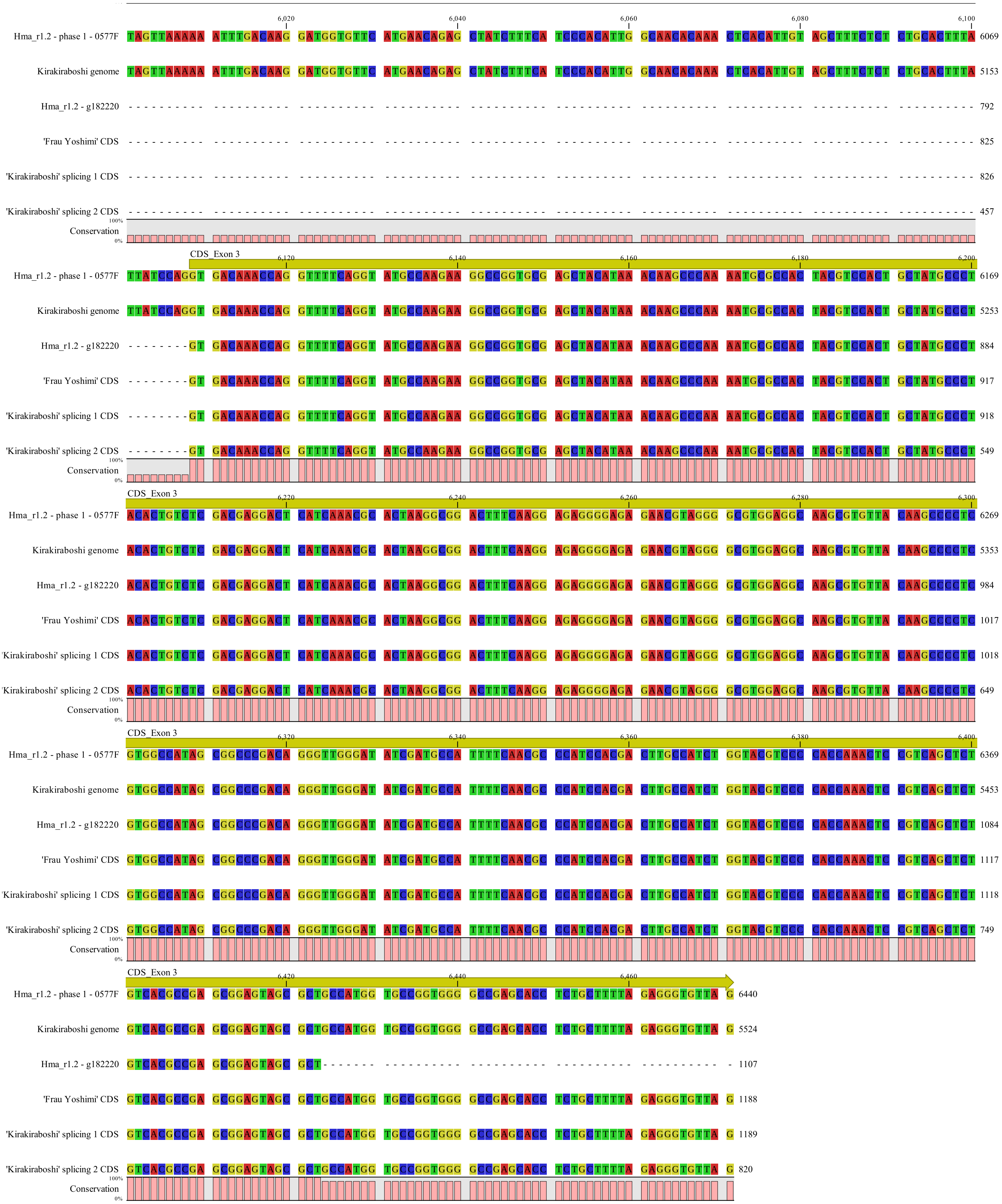


Supplementary Figure S1. Alignment of *LFY* genomic sequence and CDS. (continued)
