## Supplementary Table S1 for "Genome sequence of *Hydrangea macrophylla* and its application in analysis of the double flower phenotype"

Supplementary Table S1. RNA samples used for Iso-Seq and RNA-Seq

| Sample No. | Accession | Species | Sampled organ | Sequencing method |
|---|---|---|---|---|
| 1 | Aogashima-1 | *H. macrophylla* | Flower bud | Iso-Seq |
| 2 | Aogashima-1 | *H. macrophylla* | Flower bud | Iso-Seq |
| 3 | Aogashima-1 | *H. macrophylla* | Decorative flower | Iso-Seq |
| 4 | Aogashima-1 | *H. macrophylla* | Decorative flower | Iso-Seq |
| 5 | Aogashima-1 | *H. macrophylla* | Colored non-decorative flower | Iso-Seq |
| 6 | Aogashima-1 | *H. macrophylla* | Colorless non-decorative flower | Iso-Seq |
| 7 | Aogashima-1 | *H. macrophylla* | Fruit | Iso-Seq |
| 8 | Aogashima-1 | *H. macrophylla* | Stem | Iso-Seq |
| 9 | Aogashima-1 | *H. macrophylla* | Leaf bud | Iso-Seq |
| 10 | Aogashima-1 | *H. macrophylla* | Leaf, one-day light-intercepted | Iso-Seq |
| 11 | Aogashima-1 | *H. macrophylla* | Bud, one-day light-intercepted | Iso-Seq |
| 12 | Aogashima-1 | *H. macrophylla* | Root | Iso-Seq |
| 13 | Blue Sky | Hybrid of *H. macrophylla* and *H. serrata var. yesoensis* | Flower bud | RNA-Seq |
| 14 | Spontaneous mutant of Blue Sky | Hybrid of *H. macrophylla* and *H. serrata var. yesoensis* | Flower bud | RNA-Seq |
| 15 | S-1 | *H. macrophylla* | Flower bud | RNA-Seq |
| 16 | Spontaneous mutant of S-1 | *H. macrophylla* | Flower bud | RNA-Seq |
| 17 | Jogasaki | *H. macrophylla* | Decorative flower | RNA-Seq |
| 18 | Sumidanohanabi | *H. macrophylla* | Decorative flower | RNA-Seq |
| 19 | Sumidanohanabi | *H. macrophylla* | Non-decorative flower | RNA-Seq |
| 20 | Sumidanohanabi | *H. macrophylla* | Non-decorative flower without petal | RNA-Seq |
| 21 | Wild hydrangea 1 (collected at Niijima, Tokyo, Japan) | *H. macrophylla* | Decorative flower | RNA-Seq |
| 22 | Wild hydrangea 1 (collected at Niijima, Tokyo, Japan) | *H. macrophylla* | Non-decorative flower | RNA-Seq |
| 23 | Wild hydrangea 1 (collected at Niijima, Tokyo, Japan) | *H. macrophylla* | Flower bud | RNA-Seq |
| 24 | Wild hydrangea 2 (collected at Tateyama, Chiba, Japan) | *H. macrophylla* | Leaf | RNA-Seq |
| 25 | Wild hydrangea 3 (collected at Izu-Oshima, Tokyo, Japan) | *H. macrophylla* | Flower bud | RNA-Seq |
| 26 | Wild hydrangea 4 (collected at Shirakawa village, Gihu, Japan) | *H. serrata* | Leaf | RNA-Seq |
| 27 | Hime Ajisai | Hybrid of *H. macrophylla* and *H. serrata var. yesoensis* | Flower bud | RNA-Seq |
| 28 | Hon Ajisai | *H. macrophylla* | Flower bud | RNA-Seq |
| 29 | Mixed accessions (13, 17, 20, 21, Leaf of 'Blue Sky' and non-decorative flower of 'Jogasaki') | *H. macrophylla* and *Hybrid of H. macrophylla and H. serrata var. yesoensis* | Leaf, non-decorative flower, decorative flower and flower bud | Iso-Seq |
