## Supplementary Table S2 for "Genome sequence of *Hydrangea macrophylla* and its application in analysis of the double flower phenotype"

Supplementary Table S2 . Statistics of the genome sequences of *Hydrangea macrophylla* 'Aogashima-1'

| | HMA_r0.1 | HMA_r1.0 | | HMA_r1.1 | | HMA_r1.2 | | HMA_r1.2.pmol | |
|---|---|---|---|---|---|---|---|---|---|
| | | Primary contig | Haplotig | Phase 0 | Phase 1 | Phase 0 | Phase 1 | Phase 0 | Phase 1 |
| Total sequences | 613,685 | 3,779 | 12,012 | 3,779 | 3,779 | 3,780 | 3,780 | 18 | 18 |
| Assembly size (%) | 1,714,141,309 | 2,178,088,391 | 1,436,935,012 | 2,256,097,326 | 2,227,567,818 | 2,256,097,326 | 2,227,567,818 | 1,077,798,883 | 1,076,873,717 |
| Sequence N50 | 9,127 | 1,400,606 | 184,090 | 1,478,677 | 1,440,438 | 1,478,677 | 1,440,438 | 65,768,558 | 62,239,254 |
| Gap size (bp) | 5,985,045 | 1,051,170 | 895,550 | 1,273,060 | 1,069,590 | 1,273,060 | 1,069,590 | 673,130 | 569,310 |
| %Gap | 0.3 | >0.0 | 0.1 | 0.1 | >0.0 | 0.1 | >0.0 | 0.1 | 0.1 |
| Complete BUSCO | 72.2 | 87.3 | n.a. | n.a. | n.a. | 84.8 | 87.7 | n.a. | n.a. |
| Single-copy | 69.4 | 79.8 | n.a. | n.a. | n.a. | 76.5 | 79.0 | n.a. | n.a. |
| Duplicated | 2.8 | 7.5 | n.a. | n.a. | n.a. | 8.3 | 8.8 | n.a. | n.a. |
| Fragmented BUSCO | 12.2 | 2.7 | n.a. | n.a. | n.a. | 3.7 | 2.6 | n.a. | n.a. |
| Missing BUSCO | 15.7 | 10.0 | n.a. | n.a. | n.a. | 11.5 | 9.7 | n.a. | n.a. |
| #Genes | n.a. | n.a. | n.a. | n.a. | n.a. | 33,848 | 34,149 | n.a. | n.a. |

n.a.: not analyzed
