## Supplementary Table S3 for "Genome sequence of *Hydrangea macrophylla* and its application in analysis of the double flower phenotype"

Supplementary Table S3. J01 marker genotypes and double flower phenotypes of 15IJP1 population.

| Genotype | Phenotype | |
|---|---|---|
| | Double flower | Single flower |
| Homozygous of 117_50 allele | 39 | 0 |
| Heterozygous | 1 | 58 |
| Homozygous of 167 allele | 0 | 0 |
