## Supplementary Table S4 for "Genome sequence of *Hydrangea macrophylla* and its application in analysis of the double flower phenotype"

Supplementary Table S4. J01 marker genotypes and double flower phenotypes of 14GT77 population.

| Genotype | Phenotype | |
|---|---|---|
| | Double flower | Single flower |
| Homozygous of 117_50 allele | 18 | 0 |
| Heterozygous | 0 | 15 |
| Homozygous of 167 allele | 0 | 31 |
